# Dynamic molecular evolution of a supergene with suppressed recombination in white-throated sparrows

**DOI:** 10.1101/2022.02.28.481145

**Authors:** Hyeonsoo Jeong, Nicole M. Baran, Dan Sun, Paramita Chatterjee, Thomas S. Layman, Christopher N. Balakrishnan, Donna L. Maney, Soojin V. Yi

**Affiliations:** Georgia Institute of Technology, School of Biological Sciences, Atlanta, GA, 30332, USA; Emory University, Department of Psychology, Atlanta, GA 30322, USA; University of California, Santa Barbara, Department of Ecology, Evolution, Marine Biology, Santa Barbara, CA, 93106, USA; Karolinska Institutet, Department of Medicine Huddinge, Stockholm, Sweden; East Carolina University, Department of Biology, Greenville, NC 27858, USA

**Keywords:** supergene, evolution, selection, population genomics, gene expression, white-throated sparrow

## Abstract

In white throated sparrows, two alternative morphs differing in plumage and behavior segregate with a large chromosomal rearrangement. As with sex chromosomes such as the mammalian Y, the rearranged version of chromosome two (ZAL2^m^) is in a near-constant state of heterozygosity, offering opportunities to investigate both degenerative and selective processes during the early evolutionary stages of ‘supergenes.’ Here, we generated, synthesized, and analyzed extensive genome-scale data to better understand the forces shaping the evolution of the ZAL2^m^ chromosome in this species. We found that features of ZAL2^m^ are consistent with substantially reduced recombination and low levels of degeneration. We also found evidence that selective sweeps took place both on ZAL2^m^ and its standard counterpart, ZAL2, after the rearrangement event. Signatures of positive selection were associated with allelic bias in gene expression, suggesting that antagonistic selection has operated on gene regulation. Finally, we discovered a region exhibiting two long-range haplotypes inside the rearrangement on ZAL2^m^. These two haplotypes appear to have been maintained by balancing selection, retaining genetic diversity within the supergene. Together, our analyses illuminate mechanisms contributing to the evolution of a young chromosomal polymorphism, revealing complex selective processes acting concurrently with genetic degeneration to drive the evolution of supergenes.

## Introduction

Supergenes comprise closely linked genetic variants that are maintained due to suppressed recombination (*1, 2*). Their evolution presents an interesting paradox, in that the suppression of recombination that occurs inside supergenes reduces the efficacy of natural selection, leading to genetic degeneration. At the same time, supergenes are associated with dramatically divergent, adaptive phenotypes. These divergent phenotypes, which include classic examples of Batesian mimicry and self-incompatibility in flowering plants ((*1, 3*) and striking polymorphisms in social behavior (*4*–*10*), have long inspired both theoretical and empirical studies of their evolution. Recent genome-scale studies have illuminated wide-ranging impacts of supergene evolution on complex phenotypes across diverse taxa (e.g., (*11*–*20*)). Nevertheless, the mechanisms by which functionally divergent supergene haplotypes evolve in the face of multiple evolutionary forces remain poorly understood.

One notable example of a supergene associated with social behavior is found in white-throated sparrows (*Zonotrichia albicollis*), in which a large supergene co-segregates with parental behavior and aggression (*21*–*26*). White-throated sparrows occur in two alternative plumage morphs, white-striped and tan-striped (*27*). These morphs differ not only in their plumage coloration, but also in their social behavior, with white-striped birds exhibiting increased aggression and more frequent extra-pair copulations, and tan-striped birds engaging in more parental care compared with birds of the white-striped morph (*21, 22, 28, 29*). These alternative morphs are linked to a large (∼100 Mbp, >1k genes) rearrangement on the second largest chromosome, called ZAL2^m^, so named because the rearranged chromosome is metacentric. White-striped birds are heterozygous for ZAL2^m^ and the sub-metacentric chromosomal arrangement, ZAL2, whereas tan-striped birds are homozygous for ZAL2 (*30, 31*).

In addition to highly divergent social behavior, this relatively young supergene (estimated to have arisen 2-3 million years ago (*24, 32, 33*)), is also associated with a remarkable disassortative mating system that maintains the ‘balanced’ morph frequencies in the population. Almost all breeding pairs consist of one bird of each morph, earning the species the moniker ‘the bird with four sexes’ (*34*). Breeding pairs consisting of two individuals of the same morph are estimated to occur less than 1% of the time (*24, 30*) and only six ZAL2^m^ homozygotes (*i*.*e*., ‘super-white’ birds) have ever been identified (*23, 24, 31, 35*) out of thousands of birds karyotyped or genotyped. Given that ZAL2^m^ exists in a near-constant state of heterozygosity, it is in a state of suppressed recombination, similar to the Y and W sex chromosomes in mammals and birds, respectively. The suppression of recombination on ZAL2^m^ is expected to reduce the efficacy of natural selection, leading to reduced genetic diversity and the degeneration of the chromosome (*36, 37*). On the other hand, the tight linkage of alleles within the ZAL2^m^ supergene may contribute to adaptive phenotypes (*24, 25, 38*). Therefore, this system provides a unique opportunity to investigate the evolution of a supergene underlying social and mating behavior (*24*–*26, 39*).

Here we aim to better understand the evolutionary forces shaping the ZAL2 and ZAL2^m^ chromosomes. Our goal was to address two unanswered questions. *First, to what extent has ZAL2*^*m*^ *degenerated?* Early analyses of the rearrangement (*40*) did not show signals of degeneration, such as pseudogenization or the accumulation of repetitive sequences. However, Tuttle et al. (*24*) found a weak signal of excess non-synonymous polymorphism for genes inside the rearranged region on ZAL2^m^ and reduced allelic expression for ZAL2^m^ genes, which could be consistent with functional degradation of ZAL2^m^ (*24*). Sun et al. (*25*) similarly found a slightly higher number of non-synonymous substitutions and an increased ratio of non-synonymous to synonymous substitution rates (*d*_N_/*d*_S_) on ZAL2^m^ compared with ZAL2. Sun et al. (*25*) also found reduced expression of ZAL2^m^ alleles in brain tissue, perhaps suggesting that the accumulation of deleterious mutations has led to reduced expression of genes from ZAL2^m^. Their additional finding of reduced accumulation of mutations in functional regions suggested that ZAL2^m^ has, in fact, experienced weak purifying selection to remove deleterious alleles. Thus, while there is some evidence that ZAL2^m^ has degenerated, these results have been inconsistent and somewhat inconclusive.

### Second, what are the selective forces shaping the genomic landscapes of both ZAL2 and the ZAL2^m^ supergene?

The signals of both purifying and positive selection have been relatively weak in previous genomic analyses of ZAL2 and ZAL2^m^ (*24, 25*). Yet, by definition, ZAL2^m^ must contain variation that underlies the differences between the white-striped and tan-striped morphs (*41*–*45*). There is already some evidence that this variation affects behavior; allelic differences in the promoter region of the gene encoding estrogen-receptor alpha (*ESR1*) are likely to alter expression (*26*), and the expression of this gene was shown to be necessary for aggressive behavior typical of the white-striped morph (*26*). ZAL2^m^ is also associated with differential expression of a key neuromodulator, vasoactive intestinal peptide (*46*), known to be causal for aggression in songbirds (*47*).

Investigations of young heteromorphic sex chromosomes suggest that the accumulation of sexually antagonistic genes (*i*.*e*., genes that are beneficial to one sex and harmful to the other) may in fact drive the evolution of sex chromosomes (*48*–*50*). For example, positive selection at a small number of antagonistic alleles was shown to be a potent force shaping evolution of the young Y chromosomes in *Drosophila miranda* (*48*) even in the face of degeneration of other genes elsewhere on the chromosome. In white-throated sparrows, evidence of positive selection on both ZAL2 and ZAL2^m^ has been quite limited (*24, 25*). Nonetheless, the discovery of ZAL2- and ZAL2^m^-specific alleles that benefit the tan-striped and white-striped morphs, respectively (*26, 46*), suggests that antagonistic selection likewise contributes to the evolution of both ZAL2 and ZAL2^m^.

In addition to antagonistic selection, balancing selection may be implicated in the evolution of ZAL2^m^. The negative assortative mating system in white-throated sparrows, which maintains the chromosomal polymorphism, is a canonical example of balancing selection (*33*). However, balancing selection is also a way of maintaining advantageous genetic diversity in populations, which may be especially critical in the context of a non-recombining chromosome. Indeed, balancing selection appears to be more common in self-fertilizing (selfing) versus non-selfing species, which are likewise characterized by reduced genetic diversity, increased linkage disequilibrium, and reduced efficacy of selection (*51*–*53*). Therefore, balancing selection may maintain multiple alleles inside non-recombining regions of chromosomes like ZAL2^m^.

Previous studies have been limited in the extent to which they could test directly for degeneration, adaptive changes on ZAL2^m^, and selection at the genome level. These limitations stemmed from low sample sizes of sequencing data, the reduced intraspecies variability, and a low-quality ZAL2^m^ assembly that prevented detection of long-range haplotypes (*24, 25, 32*). Here, we overcome these challenges by analyzing extensive genomic, transcriptomic, and population data, providing insight into the evolution of young supergenes.

## Results

### Novel and extensive genomic and population data from white-throated sparrows

To better understand the evolutionary history of the ZAL2^m^ chromosomal rearrangement, we generated additional sequence data from a rare, ‘super-white’ (ZAL2^m^ homozygote) bird (*23, 25*). We generated variable fragment size libraries consisting of 150 bp paired-end reads (insert size of 300 bp and 500 bp) and 125bp mate pair reads (insert size of 1kb, 4-7kb, 7-10kb, and 10-15kb). We performed whole-genome sequencing of an additional 62 birds (49 white-striped birds and 13 tan-striped birds sampled from a variety of locations around the U.S.) (*Materials & Methods*, Supp. Table 1). White-striped birds, which are heterozygous for the rearrangement (ZAL2/2^m^), were sequenced at higher coverage than tan-striped birds (ZAL2 homozygotes) so that we could obtain sufficient reads to separate ZAL2 and ZAL2^m^ alleles in white-striped individuals (average mean depth coverages were 41.5X vs. 28.4X for white-striped and tan-striped birds, respectively, Supp. Table 2). Genomic variants were called according to the guidelines of Genome Analysis Toolkit (GATK) (ver. 4.1) (*Materials & Methods*), leading to the discovery of a total of 11,382,994 single nucleotide polymorphisms (SNPs). None of the samples showed evidence of family relationships when we computed relatedness estimates between individuals. Consequently, we used all samples in the subsequent analyses. We found a significantly higher number of polymorphic sites within white-striped birds than tan-striped birds exclusively for ZAL2/2^m^ chromosomal regions (Supp. Fig. 1). Nucleotide diversity of the ZAL2/2^m^ chromosomes was elevated in white-striped birds compared with tan-striped birds, suggesting distinctive patterns between the two plumage morphs (Fig. 1a).

**Fig. 1.**
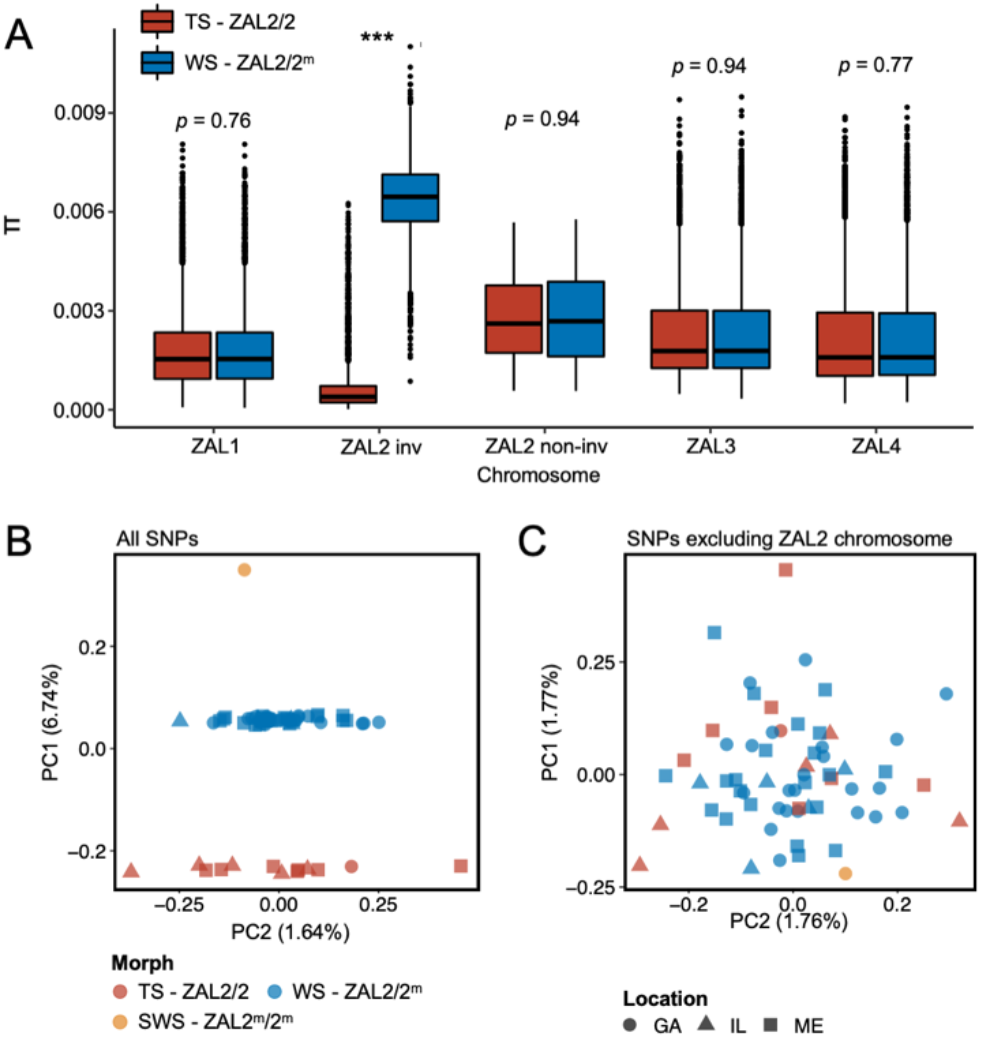
Genomic data from newly sequenced tan-striped and white-striped birds. (A) Nucleotide diversity of macro-chromosomes for tan-striped (TS) and white-striped (WS) birds. White-striped birds (ZAL2/2^m^) show elevated nucleotide diversity for the ZAL2/2^m^ inverted regions (ZAL2/2^m^ inv), while tan-striped birds (ZAL2/2) show overall reduced nucleotide diversity for the inverted regions compared with other chromosomes. (B) Scatterplots of eigenvector 1 (PC1) and eigenvector 2 (PC2) from principal component analysis of all single-nucleotide variants (SNPs) (left panel). (C) PCA excluding SNPs on the ZAL2 chromosomes (right panel). The sex chromosomes and the ZAL3 chromosome (which includes an additional chromosomal inversion) were excluded from this analysis. Note that ‘location’ here refers to the site of collection or capture of the bird. Breeding locations for GA and IL birds are unknown.

Among the total SNPs identified, 12.6% (*N* = 1,439,991) resided on scaffolds we have previously assigned to the ZAL2/2^m^ chromosome (*25*). Principal component analysis (PCA) of these ZAL2/2^m^ SNPs revealed distinct clusters corresponding to the morphs (Fig. 1b). The first principal component (PC1), which explained 6.7% of the variation in the data, clearly separated tan-striped and white-striped birds, with the lone super-white individual (ZAL2^m^/2^m^ homozygote) as a clear outlier. In contrast, other available phenotypic information, including sex and geographic origin of samples, did not show meaningful variation with the principal components, and other PCs had little explanatory power (Fig. 1b). Tests for admixture also failed to identify significant population substructures by geographical origin of samples (Supp. Fig. 2). This lack of population structure is unsurprising, as 35 of the 63 samples (56%) were from birds that were migrating, and, thus, the breeding location of these birds is unknown.

### Features of the ZAL2^m^ chromosome consistent with reduced efficacy of natural selection and low levels of recombination

We examined several genomic features of the ZAL2^m^ chromosome using the additional genomic resources we generated. We first performed a *de novo* genome assembly of the super-white bird, employing newly generated sequence data, to study the ZAL2^m^ chromosome with an assembly derived entirely from a bird homozygous for the ZAL2^m^ chromosome. The total assembly size was 1,058 Mbp (N50 length of 3.1 Mbp, longest scaffold 27 Mbps), comparable to that of the ZAL2/2 reference assembly (1,052 Mbp, N50 scaffold length of 4.86 Mbp, longest scaffold 45 Mbp) (see Supp. Table 3 for more details). There were 160 putatively ZAL2^m^-linked scaffolds (*Materials & Methods*), with a total length (110.99 Mbp) comparable with that of ZAL2-linked scaffolds from the reference assembly (108.5 Mbp (*24*)). Despite this similarity in total length, however, the average length of the individual ZAL2^m^-linked scaffolds was significantly shorter than scaffolds on other chromosomes in the super-white assembly (*p* < 0.001, Mann-Whitney U-test). It was also shorter than the average scaffold length on the ZAL2 chromosome in the ZAL2/2 reference assembly (Fig. 2a). We did not observe such a pattern in the other chromosomes of similar size when comparing between the two assemblies (Fig. 2a). This result was consistent with the presence of repetitive DNA sequences on ZAL2^m^ causing more assembly breaks compared with the ZAL2/2 reference genome. We found evidence that the ZAL2^m^ chromosome contained more repeat elements and was especially enriched for long terminal repeat (LTR) elements (2.4 Mbp vs. 2.1 Mbp) and interspersed repeats (5.8 Mbp vs. 5.5 Mbp), compared with the ZAL2 chromosome. The number of these repeat elements is likely to be underestimated, given that the ZAL2^m^ assembly is highly fragmented. Additionally, we found that ZAL2 and ZAL2^m^ had accumulated a higher proportion of structural variants (insertions and deletions) compared with other chromosomes (Fig. 2b).

**Fig. 2.**
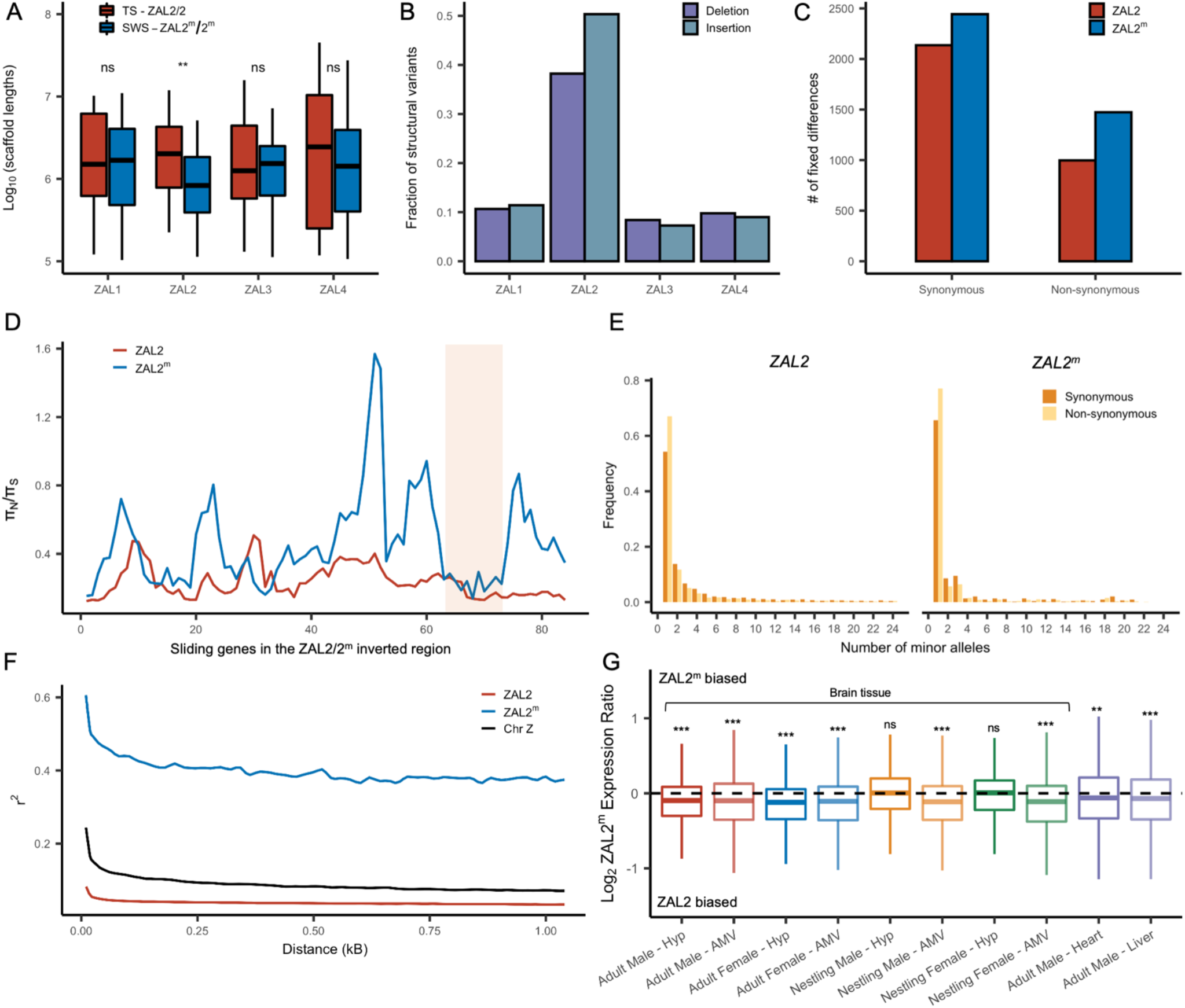
Genetic divergence between ZAL2 and ZAL2^m^ chromosomes. (A) The scaffolds for the ZAL2^m^ chromosome in the super-white (SWS) assembly tend to be fragmented compared with those for the ZAL2 chromosome in the tan-striped (TS) assembly. ** *p* < 0.001 (Mann-Whitney U-test); ns, not significant (B) Fraction of structural variants (SV), both insertion and deletion events, for the 4 largest chromosomes, using the tan-striped assembly as a reference. The fraction of SV is computed as a total base affected by variants divided by the length of chromosome. (C) Number of fixed protein-coding mutation derived in ZAL2 and ZAL2^m^. (D) Sliding window (window size of 20 genes with step size of 5 genes) analysis of the ratio of nonsynonymous to synonymous nucleotide diversity (π_N_/π_S_) within the ZAL2 and ZAL2^m^ chromosomes. The ZAL2^m^ outlier region is highlighted (colored background). (E) Site frequency spectrum of polymorphic sites. (F) Decay of linkage disequilibrium. (G) Proportion of the ZAL2^m^ allele expressed for each tissue set. The proportion of the ZAL2^m^ allele expressed is less than the null hypothesis of 0.5 for all tissues except nestling AMV using FDR correction. Hyp, hypothalamus; AMV, ventromedial arcopallium.

Next, using the large amount of newly generated population genomic data, we examined patterns of SNPs on ZAL2^m^ alleles separately from those on ZAL2 via haplotype phasing using fixed differences between the two chromosome types (*Materials & Methods*). We found that the total number of genetic variants was approximately 12-fold reduced on the ZAL2^m^ alleles compared with ZAL2 alleles (29,420 vs. 367,466 SNPs and 3,921 vs. 32,479 indels on ZAL2^m^ and ZAL2, respectively, after excluding singleton variants). The mean nucleotide diversity was similarly reduced on the ZAL2^m^ chromosome compared with the ZAL2 chromosome (0.0104% vs. 0.1141% for ZAL2^m^ vs. ZAL2, respectively).

Analyses of the genetic variants on the ZAL2 and ZAL2^m^ alleles showed evidence of only weak genetic degeneration. The ratio of non-synonymous to synonymous fixed differences inside the rearranged region was slightly, but significantly, elevated for ZAL2^m^-derived compared with ZAL2-derived fixed differences (Fig. 2c), which is consistent with either positive selection or inefficient purifying selection on ZAL2^m^. We found that the ratio of non-synonymous to synonymous nucleotide diversity (π_N_/π_S_) was significantly increased on ZAL2^m^ compared with ZAL2 (*p* = 1.3 × 10^−13^, Mann-Whitney U-test, Fig. 2d). The minor allele site frequency spectrum (SFS) for the ZAL2^m^ synonymous and non-synonymous sites showed a large proportion of singleton variants and an irregular decay of allele frequency as the minor allele count increases (Fig. 2e), also suggesting reduced efficacy of purifying selection on ZAL2^m^.

Assuming the mutation rates of the two chromosomes are similar, the ratio of effective population size (*N*_e_) between the ZAL2^m^ and ZAL2 can be approximated by the ratio of nucleotide diversity of synonymous sites between the ZAL2 and ZAL2^m^. The proportion of *N*_e_ between ZAL2^m^ and ZAL2 is 0.12 (± 0.01), which is 3-fold lower than the expected ratio of 0.33 (because the ZAL2^m^ chromosome is 1/3 as frequent as the ZAL2). This lower proportion suggests that *N*_e_ of ZAL2^m^ has undergone further reduction than expected from the census size in the population, consistent with the effects of reduced recombination. We noted that the linkage disequilibrium between variants on ZAL2^m^ exhibited the classic decay with distance (Fig. 2f), indicating at least some level of recombination. Taken together, these results are consistent with reduced, but not entirely eliminated, recombination on the ZAL2^m^ chromosome.

We next examined whether degeneration inside the rearranged region on ZAL2^m^ has resulted in globally reduced expression of the ZAL2^m^ allele. To do so, we used multiple large RNAseq datasets from a variety of tissues in birds sampled from different geographic locations and times of year (see *Materials & Methods*, Table 1). As predicted and consistent with what was previously reported (*25*), we found evidence of consistently reduced expression of the ZAL2^m^ allele in 8/10 types of tissue (Fig. 2g). We next tested for an association between the number of accumulated mutations (non-synonymous, synonymous, and in the promoter region) on ZAL2^m^ and allelic bias (AB) in expression of the ZAL2^m^ allele within each tissue, which would link genetic degeneration within or near genes to reduced expression of ZAL2^m^. We found evidence that allelic bias in gene expression was associated with the decile rank of the per-base number of non-synonymous fixed differences (*X*^2^(1) = 12.54, *p* = 0.000398), although the effect size was exceedingly small (marginal *r*^2^ = 0.0026). Neither the decile rank of the per-base number of synonymous fixed differences (*X*^2^(1) = 0.9335, *p* = 0.334) nor the number of mutations within 1kb upstream of the transcription start site (*X*^2^(1) = 0.8992, *p* = 0.343) were associated with allelic bias. Thus, the overall reduction in expression of the ZAL2^m^ allele is weakly associated with an increased number of non-synonymous fixed nucleotide changes within genes. The limited and weak nature of the effect suggests, however, that the pattern of gene expression may have been affected also by other factors, for example ongoing selection (thus manifested as nucleotide polymorphism), selection at more distal sites, and/or epigenomic mechanisms, such as differences between ZAL2 and ZAL2^m^ in DNA methylation or histone modification (see (*39*)).

**Table 1.** List of RNA sequencing data sets

|  | Tissue | Sample Size (WS/total) | Collection details | Source |
| --- | --- | --- | --- | --- |
| Adult males | Brain (Hyp, AMV) | 9/20 | Collected early in the breeding season | Zinzow-Kramer et al, 2015; Sun et al., 2018<br>Accession: GSE77186 |
| Adult females | Brain (Hyp, AMV) | 6/11 | Collected early in the breeding season | Accession: PRJNA657006 |
| Nestlings (both sexes) | Brain (Hyp, AMV) | 16/32 | Collected from nests during the breeding season | Accession: PRJNA657006 |
| Adult males (all white-striped ) | Heart & Liver | 20/20 | Collected during fall migration, then housed in captivity on either long or short days to simulate breeding vs. non-breeding; collected at two time points during the day | Horton et al., 2019<br>Accession: GSE116989 |

### Evidence of regional balancing selection on the ZAL2^m^ chromosome

Although the level of genetic diversity was overall reduced on ZAL2^m^, it was exceptionally elevated in one region corresponding to ∼3Mbps spanning 5 scaffolds. This region, henceforth referred to as the ZAL2^m^ outlier region (Fig. 2d, 3a), includes at least 15 protein-coding genes that are well conserved as single copy genes across 13 avian species (Table 2). On average, nucleotide diversity in ZAL2^m^ across this region was 0.001, which is 10-fold higher than the mean nucleotide diversity of ZAL2^m^ and even exceeds the nucleotide diversity in the corresponding region within ZAL2 by 3.15-fold.

**Table 2.** List of protein-coding genes inside the ZAL2m outlier region

| Gene | Scaffold | Start | End | $\pi$ ZAL2 | $\pi$ ZAL2<sup>m</sup> | TaD_ZAL2 | TaD_ZAL2<sup>m</sup> | $D_{XY}$ |
| --- | --- | --- | --- | --- | --- | --- | --- | --- |
| KCNS3 | NW_005081621.1 | 97089 | 110512 | 2.75E-04 | 8.43E-04 | -1.2953 | -0.8658 | 0.011281 |
| MSGN1 | NW_005081621.1 | 160375 | 160897 | NA | 6.10E-04 | NA | -0.1138 | 0.003056 |
| GEN1 | NW_005081621.1 | 175791 | 198245 | 3.52E-04 | 2.22E-03 | -1.1302 | 1.1728 | 0.011647 |
| SMC6 | NW_005081621.1 | 198452 | 244136 | 3.94E-04 | 2.27E-03 | -1.0088 | 0.8548 | 0.011901 |
| MYCN | NW_005081621.1 | 1179492 | 1184761 | 2.59E-04 | 6.18E-04 | -0.5898 | 0.2116 | 0.005173 |
| DDX1 | NW_005081621.1 | 1432697 | 1452535 | 3.41E-04 | 2.42E-03 | -1.5819 | 0.8752 | 0.014699 |
| NBAS | NW_005081621.1 | 1454601 | 1615580 | 2.93E-04 | 2.17E-03 | -1.6271 | 2.2291 | 0.012296 |
| TRIB2 | NW_005081621.1 | 2596178 | 2616950 | 3.45E-04 | 1.89E-04 | -1.0009 | -0.8376 | 0.011498 |
| LPIN1 | NW_005081621.1 | 3012153 | 3061203 | 2.73E-04 | 3.27E-04 | -1.7799 | -0.7252 | 0.015295 |
| GREB1 | NW_005081621.1 | 3100814 | 3165186 | 2.25E-04 | 1.72E-04 | -1.8299 | -0.7351 | 0.01458 |
| E2F6 | NW_005081582.1 | 24475 | 46577 | 1.90E-04 | 1.59E-04 | -1.5978 | -1.0219 | 1.4E-02 |
| ROCK2 | NW_005081582.1 | 50993 | 155170 | 2.62E-04 | 6.34E-04 | -1.46 | -0.2362 | 1.4E-02 |
| KCNF1 | NW_005081582.1 | 331424 | 333991 | 9.85E-05 | 2.41E-04 | -0.2519 | -0.8641 | 5.9E-03 |
| PDI A6 | NW_005081582.1 | 415853 | 431919 | 3.74E-04 | 1.49E-04 | -1.1579 | -1.2521 | 1.2E-02 |
| ATP6V1C2 | NW_005081582.1 | 431766 | 454886 | 2.56E-04 | 6.94E-04 | -1.7535 | -0.2391 | 1.4E-02 |

To first examine whether this region has recently experienced an introgression event that could have caused the observed patterns, we constructed a phylogenetic tree of this region. The resulting tree exhibited the same topology as those from other regions of the ZAL2^m^, thus providing no support for introgression (Fig. 3b). In addition, we calculated the D-statistic (ABBA-BABA test (*53*)) to examine patterns of introgression with other species from the same genus, *Zonotrichia querula* and *Zonotrichia atricapilla*. Sliding window estimates of the D-statistic (ABBA-BABA statistic) did not show any differences in patterns between haplotypes (Supp. Fig. 4), suggesting that the multiple haplotype structures in ZAL2^m^ were not introduced by introgression. Nonetheless, the ZAL2^m^ outlier region showed longer branch lengths than did both the corresponding region on the ZAL2 chromosome and a randomly selected region of ZAL2^m^, reflecting the high genetic diversity within this region (Fig. 3b). Tajima’s D was significantly higher throughout the ZAL2^m^ outlier region compared with the genomic background (*p* < 0.001, permutation test) (Fig. 3a).

**Fig. 3.**
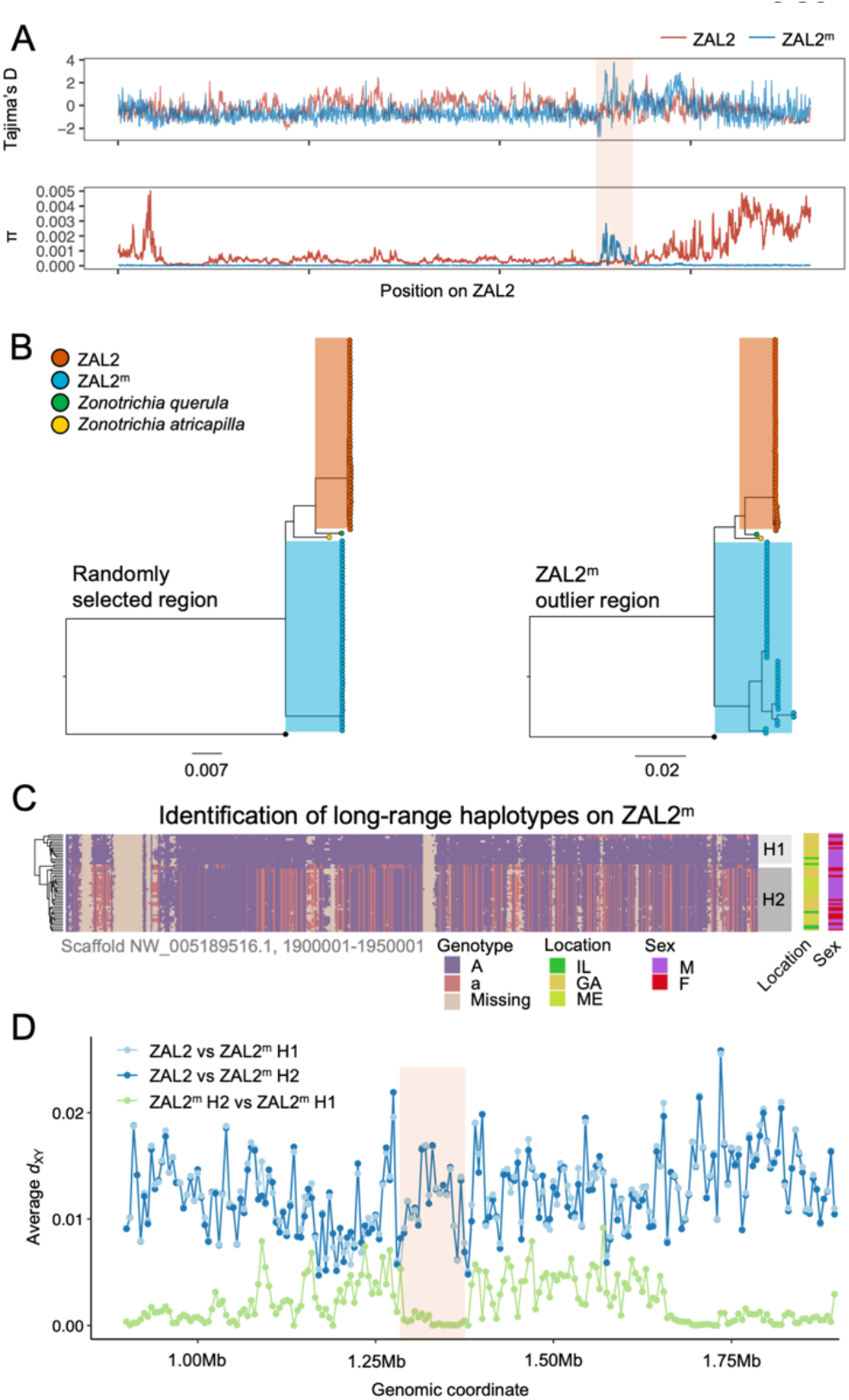
Genetic diversity and patterns of divergence across the rearranged region of the ZAL2^m^ chromosome and in the ZAL2^m^ outlier region. (A) Tajima’s D and nucleotide diversity across the ZAL2 and ZAL2^m^ chromosomes. The ZAL2^m^ outlier region is highlighted (colored background). (B) Phylogenetic tree of randomly selected regions (left panel) and the ZAL2^m^ outlier region (right panel). The ZAL2^m^ chromosome shows multiple haplotype structures and has longer branch lengths within the population compared with ZAL2 chromosomes. (C) Single nucleotide polymorphism genotype plot of a scaffold inside the ZAL2^m^ outlier region (Scaffold NW_005189516.1, 1900001-1950001). The plot shows two haplogroups. Major allele SNPs (A, same genotype as the super-white ZAL2^m^/2^m^ genome) are represented in purple, minor allele (a, different from the super-white genome) in red, and missing data in tan. (D) Genetic divergence (*d*_XY_) for a portion of the rearrangement. *d*_XY_ between the ZAL2 chromosome and haplogroup 1 (H1) is plotted in light blue, between ZAL2 and haplogroup 2 (H2) in dark blue, and between H1 and H2 in light green.

The increase of nucleotide diversity, high Tajima’s D, and long branch length all suggest that balancing selection has impacted the ZAL2^m^ outlier region. Concordantly, we identified a signature of balancing selection in the ZAL2^m^ outlier region on the basis of allele frequency spectrum (*β* statistics) computed using BetaScan (*54*). Visual examinations of the ZAL2^m^ outlier region also revealed the presence of two long-range haplotypes (tentatively called haplogroup 1 and haplogroup 2) that likely reflect long-term balancing selection and several recombination events within the haplotypes (Fig. 3c). The genetic differentiation between the two ZAL2^m^ outlier haplogroups was lower than the differentiation between either haplogroup and ZAL2 (Fig. 3d), indicating that the haplogroups have evolved since the split between ZAL2 and ZAL2^m^. This interpretation is also consistent with the inferred phylogeny (Fig. 3b). The ZAL2^m^ outlier regions exhibited an excess of non-synonymous SNPs compared with synonymous SNPs (Figure 2d), as well as an excess of intermediate frequency minor alleles (Supp. Fig. 5). Neither geographic location of collection nor sex was associated with any distinct patterns by haplogroups in a PCA analysis using genetic variants in the region (Supp. Fig. 6; although note the breeding location of the birds from two of the locations was unknown). Together, these results solidly implicate balancing selection in the evolution of the ZAL2^m^ outlier region.

We looked for phenotypic consequences of the putative ZAL2^m^ haplogroups using the subset of white-striped birds from the sequencing samples for which we had phenotypic data (*N* = 6 for haplogroup 1, *N* = 12 for haplogroup 2, see (*55, 56*) for details). We did not find an effect of haplogroup on aggressive behavior (measured during simulated territorial intrusions), gonad size, or cloacal protuberance volume. We found a trend for tarsus length, a measure of skeletal frame size (Supp. Fig. 7), although this effect was not significant when correcting for multiple testing.

We next tested for effects of haplogroup on the expression of genes inside the ZAL2^m^ outlier region. In the subset of white-striped birds for which we had haplogroup information, we examined RNAseq data from two brain regions, the hypothalamus (Hyp) and the ventromedial arcopallium (AMV, the functional homolog of the medial amygdala, formally named the nucleus taeniae of the amygdala) (see *Materials & Methods* and Supp. Table 4 for details). We found that the gene *GREB1* (Growth Regulating Estrogen Receptor Binding 1) was more highly expressed in haplogroup 2 (unadjusted *p* = 0.0195) in AMV. Additionally, there was a trend for the gene *KCNS3* to be more highly expressed in both Hyp (unadjusted *p* = 0.0511) and AMV (unadjusted *p* = 0.1567). This gene encodes a voltage-gated channel subunit that in humans and mice is specific to fast-spiking parvalbumin (inhibitory) neurons (*57, 58*). These findings provide potentially interesting candidates to explore further in the context of behavioral differences between the morphs.

### Candidates for positive selection in ZAL2 and ZAL2^m^

Our work has provided a refined map of genetic differentiation between the ZAL2 and ZAL2^m^ chromosomes (Supp. Fig. 8). Interestingly, three scaffolds near the chromosomal end (*i*.*e*., at the beginning of the p-arm, as well as at the end of the q-arm in the genomic coordinate of the ZAL2 chromosome), exhibited both decreased *F*_*ST*_ values and a reduced rate of fixed genetic differences (D_f_). One possible explanation for these findings is that these regions represent a younger evolutionary stratum (*59*–*61*). Another possibility is that the within-ZAL2 nucleotide diversity is increased and that this inflated diversity within the sampled population reduced the *F*_*ST*_ estimate.

By leveraging our resource of a haplotype map of both the ZAL2 and ZAL2^m^ chromosomes, we investigated signatures of positive selection in each chromosome. Based on the H-statistics using the lengths of linkage disequilibrium between haplotypes ((*62*), *Materials & Methods*), we identified 216 regions in ZAL2 exhibiting a significantly elevated H-statistic (empirical *p*-values < 0.05). Notably, of these regions, four top candidate regions (NW_005081582.1, 480-520kb and 920-960kb) showed evidence of positive selection on both the ZAL2 and ZAL2^m^ chromosomes (Supp. Fig. 9) (*32*). We found significant peaks also in the region corresponding to the chromosome end in ZAL2 (Supp. Fig. 9). These observations suggest that selective sweeps took place on both ZAL2 and ZAL2^m^ chromosomes after the chromosomal rearrangement events. We also identified a long stretch (6 Mbp region, 365-650 windows in Supp. Fig. 9a) showing an overall elevated H-statistic in ZAL2. This region also exhibited the lowest estimates of nucleotide diversity within ZAL2 and the highest estimates of *F*_*ST*_ between the two chromosome types. There are 68 genes located in regions that have a significant (p < 0.05) H-statistic for ZAL2^m^ and 109 genes in regions with a significant H-statistic for ZAL2 (Supp. Table 5), meaning that they are inside regions showing evidence of a selective sweep, but we found no evidence of ontological or functional enrichment for these genes.

### Antagonistic selection influences ZAL2^m^ gene expression

Some of the behaviors that differ between morphs, namely territorial aggression, are predicted by the expression of particular candidate genes inside the ZAL2/2^m^ rearrangement (*26, 38*). Because the rearrangement determines morph, we hypothesized that the evolution of ZAL2 and ZAL2^m^ was facilitated by antagonistic selection for alleles that benefit individuals of the tan-striped and white-striped morph, respectively. To test this antagonistic selection hypothesis, we examined whether allelic bias in expression of genes inside the rearrangement was associated with differential expression of those genes between the tan-striped and white-striped morphs (lists of genes showing differential expression or allelic bias can be found in Supp. Table 6). Allelic bias in expression, in this case differential expression between the ZAL2 and ZAL2^m^ alleles in white striped birds (see Fig. 2g), would indicate possible effects of *cis*-regulatory variants. For these tests, we used data on gene expression in the brain only, as heart and liver tissue samples were limited to white-striped birds.

First, we found that among the genes that were differentially expressed between morphs, more of these genes are located inside the rearranged region on ZAL2/2^m^ than expected by chance (*X*^2^ tests showed highly significant effects for both brain regions in adults and nestlings of both sexes, with FDR correction) (Fig. 4a). Next, we aimed to test whether the differentially expressed genes showed greater allelic bias than genes without differential expression. Consistent with what we previously reported using only the males (*25*), we found that differential expression significantly predicted the degree of allelic bias in expression of that gene (*X*^2^(2) = 664.16, *p* < 2.2 × 10^−16^, controlling for sequencing batch and brain region, see *Materials & Methods*). In addition, tan-biased genes showed greater ZAL2 allelic bias and white-biased genes showed greater ZAL2^m^ allelic bias than did genes that were not differentially expressed between morphs (post-hoc T>W vs. T=W: *z* = -16.87; post-hoc W>T vs. T=W: *z* = 19.63, *p* < *2*.*2* × 10^−16^; post-hoc W>T vs. T>W: *z* = 26.35, *p* < 2.2 × 10^−16^) (Fig. 4b). These findings suggest that differences in gene expression between the morphs are driven, in part, by evolutionary changes in the regulatory regions of genes captured by the rearrangement.

**Fig. 4.**
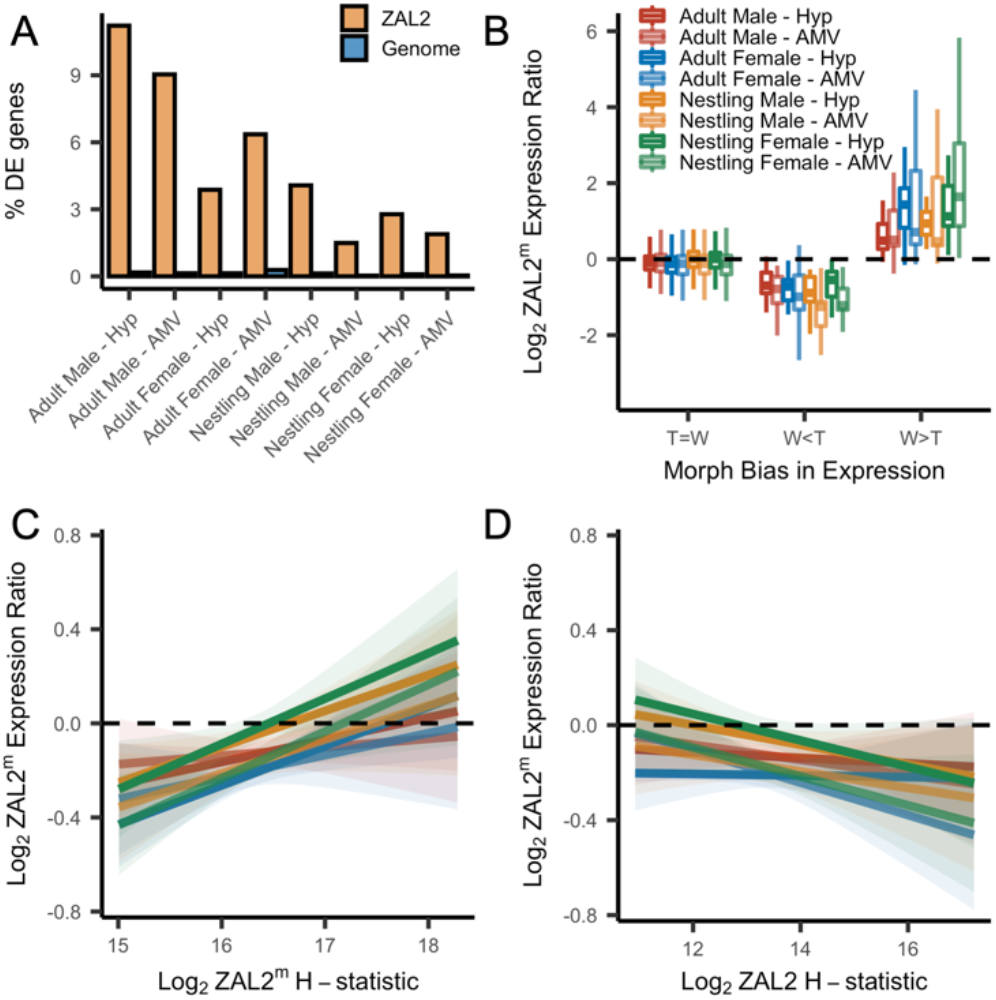
Evidence for antagonistic selection driving ZAL2 and ZAL2^m^ gene expression in the brain. (A) shows the percentage of differentially expressed genes that reside inside the rearranged region on ZAL2, versus elsewhere in the genome. The percentage of differentially expressed genes inside vs. outside the rearranged region of ZAL2 is higher than expected by chance (*p*_adj_ < 2.2 × 10^−16^ for all comparisons). (B) shows log_2_ ZAL2^m^ expression ratios for genes that were more highly expressed in white-striped birds (W>T), genes more highly expressed in tan-striped birds (T>W) and those that that do not significantly differ between morphs (T=W). (C) Log_2_ ZAL2^m^ expression ratios are plotted versus the Log_2_ ZAL2^m^ H-statistic for each category of sample. Hypothalamus (Hyp), Ventromedial arcopallium (AMV). (D) Log_2_ ZAL2^m^ expression ratio are plotted versus the Log_2_ ZAL2 H-statistic.

We also looked for a relationship between gene expression and signatures of positive selection on both the ZAL2 and ZAL2^m^ chromosomes. We therefore asked whether genes located in regions with evidence for a selective sweep on either ZAL2 or ZAL2^m^ (as indicated by the H-statistics for 50kb windows) were also likely to exhibit bias in gene expression in the brain. For the ZAL2^m^ chromosome, we found that the ZAL2^m^ H-statistic was a significant, positive predictor of bias in brain gene expression towards the ZAL2^m^ allele, controlling for sequencing batch and brain region (*X*^2^(2) = 40.17, *p* = 1.893 × 10^−9^; *t* = 5.114, *p* = 3.24 × 10^−7^) (see Materials & Methods) (Fig. 4c). Likewise, we found that genes that exhibited evidence of positive selection on ZAL2 showed allelic bias towards the ZAL2 allele (*X*^2^(2) = 24.231, *p* = 5.475 × 10^−6^; *t* = -3.151, *p* = 0.00164) (Fig. 4d). Taken together, these results suggest that gene expression on ZAL2 and ZAL2^m^ is driven by selection for alleles that benefit each morph.

## Discussion

Understanding evolutionary processes that contribute to the origin and maintenance of supergenes can elucidate the links between genes and complex phenotypes such as behavior. One of the most critical processes in the evolution of supergenes is the suppression of recombination (*1*), a consequence of which is genetic degeneration. Such processes have been extensively studied in the case of non-recombining sex chromosomes (*36, 37, 63, 64*), but their role in the early evolution of non-recombining autosomes is not well understood. The ZAL2^m^ supergene captures a snapshot of an early stage of supergene evolution. We and others have previously observed weak signatures of degeneration on the ZAL2^m^ chromosome (*24, 25*). In this study, using a large, newly generated genomic sequencing and population genomic data set, we demonstrate pervasive signatures of genetic degeneration at multiple levels, from genomic contigs to SNP density. Regardless, the degeneration is weak at most, and it appears that recombination has not entirely ceased on ZAL2^m^. It is likely that this low level of recombination occurs in rare ZAL2^m^ homozygotes (*23*). Therefore, the case of the ZAL2^m^ supergene illustrates that complex phenotypes, including alternative plumage morphs and life history strategies, can evolve prior to the cessation of recombination and despite substantial genetic degeneration.

As is evident from research on non-recombining sex chromosomes, both positive and antagonistic selection can further the differentiation between those chromosomes (*47, 65*–*69*). Previous studies of the ZAL2^m^ supergene revealed no or only weak evidence of positive selection at the molecular level (*24, 25*), despite evidence that allelic biases in expression may contribute causally to phenotypic differences between white-striped and tan-striped morphs (*26*). Here, using extensive molecular data, we detected a strong signature of antagonistic selection based on gene expression profiles and identified several regions of positive selection on both ZAL2 and ZAL2^m^. We also found that allele-specific gene expression in the brain was associated with these signatures of positive selection on both the ZAL2 and ZAL2^m^ chromosomes, suggesting that this selection has acted on regulatory regions of genes. Several regions showed evidence of selective sweeps, including regions near the ends of the chromosomes, which is worth noting given that positive selection near inversion breakpoints has been found in other species including *Drosophila* (*70*). Furthermore, we observed signatures of positive selection on large regions of ZAL2, which was reminiscent of the finding that the X chromosome of primates has been targeted by strong positive selection (*71*). This positive selection on ZAL2 could indicate that recessive alleles on that chromosome experience increased selection in birds of the white-striped morph, and are thus in direct conflict with genes on ZAL2^m^.

Remarkably, we also discovered clear signatures of balancing selection maintaining two long-range haplotypes within the ZAL2^m^ supergene. We were surprised by this finding, as similar long-range haplotypes have not, to our knowledge, been previously reported for non-recombining chromosomal polymorphisms. It has been speculated that balancing selection is less common in sexually reproducing species than in self-fertilizing species (*50*), in which the effective recombination rate is lowered due to the extreme degree of inbreeding. Our results suggest that balancing selection may function to maintain genetic variability in the face of reduced recombination, and we might expect to see other instances of balancing selection within supergenes.

The phenotypic effects of these balanced haplotypes remain unknown. We found some evidence of an association between haplotype and a measure of body size, although this relationship needs to be investigated further with a greater number of individuals. In addition, two out of the fifteen genes inside the balanced ZAL2^m^ outlier region showed signatures of differential gene expression between the haplogroups. One of these genes is *GREB1*, which in mammals is a downstream target of estrogen receptor alpha (ERα) (*72*). This finding is notable because in white-throated sparrows, ERα is causally related to the behavioral differences between morphs (*26, 56*). Thus, the ZAL2^m^ haplogroup may interact in important ways with ERα-mediated phenotypes in white-striped birds, although further research is needed to test this hypothesis.

Taken together, our results demonstrate that the ZAL2^m^ supergene is far from a degenerating counterpart of a ‘normal’ autosome. Rather, it is an active arena of multiple evolutionary forces. Although most studies of the consequences of suppressed recombination have focused on genetic degeneration, the resulting supergenes also give rise to opportunities for evolutionary innovation (*1, 2, 11*). Our results show that although degeneration is operating on ZAL2^m^, dynamic selective forces are occurring simultaneously, producing divergent complex phenotypes. These results emphasize that supergenes are an important force in adaptive evolution (*1, 11, 73*) and open the door for future studies to demonstrate and investigate the functional consequences of dynamic natural selection acting inside recombination-suppressed supergenes.

## Materials & Methods

### Whole genome sequencing and population genetics

We performed whole genome sequencing of 63 samples (WS: *N* = 49, TS: *N* = 13, and SW: *N* =1), which included 28 birds captured during the breeding season near Argyle, ME, USA, 25 birds captured in Atlanta, GA, USA during fall migration (November & December), and 10 post-mortem samples opportunistically collected from birds who died from building strikes in Chicago, IL, USA (see (*66*)) during the spring or fall migrations. Sequencing reads from these samples, as well as reads from three previously described samples, were mapped to the reference genome assembly (GCF_000385455.1) using Bowtie2 (ver. 2.3.5) with the –very-sensitive-local option. Possible PCR duplicates were removed using Picard tools (ver. 2.19). SNP and INDEL calling were conducted using GATK Haplotypecaller (ver. 4.1.2) with the -ERC GVCF option and joint genotyping of all samples were performed with GenotypeGVCF option. We filtered out SNPs with any missing information, MAF <0.05, meanDP <5, or meanDP>80. Raw sequencing reads are available on NCBI SRA (BioProject PRJNA818012).

### Sequencing and genome assembly of super-white bird

To complement the available short-read sequencing data from the female super-white bird (*25*), we generated additional paired-end reads from the same individual (150X; insert size of 300 bp and 500 bp) to improve the assembly quality. Genomic DNA was extracted from a 200mg liver sample from the super-white bird using the Qiagen DNEasy Blood and Tissue DNA kit. Additionally, mate-pair libraries of different insert sizes (insert size of 1kb, 4-7kb, 7-10kb, and 10-15kb) were prepared and sequenced by the Brigham Young University Genome Sequencing Center. Raw sequencing reads are available on NCBI SRA (BioProject PRJNA818012). Using these data, we constructed a whole genome *de novo* assembly of *Z. albicollis*, including the ZAL2^m^ chromosome. Paired-end sequencing reads were trimmed by Trimmomatic v.0.32 (*76*), and error correction of the trimmed sequencing reads was conducted by Lighter v.1.1.2 (*77*). Initial contig assembly and scaffolding was conducted by ABySS v.2.1.5 (*78*). We used Gapcloser (*79*) to fill the gaps emerging from scaffolding process. The total assembly size was 1,058 Mbp (N50 length of 3.1 Mbp), consisting of 160 scaffolds (111.76 Mbp) belonging to the ZAL2^m^ chromosome with the longest scaffold length of 5.1 Mbp (Supp. Table 3).

### Identification of ZAL2/2^m^ scaffolds

To discriminate scaffolds that originate from the ZAL2 versus the ZAL2^m^ chromosome, we mapped the super-white assembly against the genome of the House sparrow (*Passer domesticus*), which is the most closely related species with chromosome level assembly using LASTZ v1.03. Scaffolds uniquely mapped to the homologous House sparrow chromosomes with >30% coverage and >85% identity were retained. In the case of multi-mapping scaffolds, we used more stringent criteria with >70% coverage and >85% identity. We conducted the same procedure using the tan-striped reference assembly. The list of matched scaffolds was highly consistent with our previous study, with the addition of several new scaffolds (N = 15, sum of scaffold lengths = 129.1kb). To distinguish sequences inside vs. outside of the rearranged (*i*.*e*., inverted) regions, we computed the average frequency of heterozygous SNPs in sliding windows of 25kb size with 1kb step.

### Genetic differentiation between the ZAL2 and ZAL2^m^ chromosomes

Utilizing the large amount of newly generated whole genome sequence data, here we identified putatively fixed genetic differences between the ZAL2 and ZAL2^m^ chromosomes. Briefly, we identified positions at which the genomes of the tan-striped (ZAL2/2) and super-white (ZAL2^m^/2^m^) birds are homozygous for different alleles while the white-striped (ZAL2/2^m^) birds are heterozygous. Note that the probability of this allelic pattern occurring by random chance is 2.6 × 10^−23^ given the sample size, according to a binomial test. Following this procedure, we obtained a total of 931,424 SNPs and 97,375 insertions and deletions (InDels) between ZAL2 and ZAL2^m^ chromosomes, increasing the number of putatively fixed SNPs between the two chromosomes beyond previous publications (*24, 25*). Variant call format files of fixed differences (both SNPs and InDels) are available at DOI: 10.6084/m9.figshare.19395146. As we expected, a vast majority (N = 930,588; 99.91%) of the fixed differences reside in the scaffolds that we previously predicted to be inside the rearrangement (*25*). The remaining fixed differences were found in scaffolds that were either too short or that mapped ambiguously to multiple chromosomes.

### Haplotype phasing of whole genome and RNA sequencing data

We performed read-based haplotype phasing of the sequencing data using reads from white-striped individuals (ZAL2/2^m^ heterozygotes). Briefly, using the 931,424 single nucleotide fixed differences between ZAL2/2^m^, we assigned sequence reads from the heterozygous birds to the corresponding chromosome of origin (*i*.*e*., either ZAL2 or ZAL2^m^) if the paired-end reads were mapped to a region that overlapped at least one fixed difference. Reads from white-striped birds were mapped to *N*-masked (masking the putative fixed differences between ZAL2 and ZAL2^m^) reference assembly to avoid mapping bias and assigned to their chromosome of origin using SNPsplit v.0.3.2. On average of 8.5% of all reads from white-striped birds were assigned to either the ZAL2 or ZAL2^m^ chromosomes. In comparison, the size of the chromosomal inversion is estimated to be approximately 9.5% of the total genome (*32*). The ZAL2 and ZAL2^m^ assigned reads were extracted respectively using Bedtools *bamtofastq*. To call variants for both ZAL2 and ZAL2^m^ chromosomes, the reads were re-mapped to the reference genome assembly using Bowtie2 (ver. 2.3.5). Variant calling was conducted using GATK *Haplotypecaller* (ver. 4.1.2) with the -ERC GVCF option. Vcftools was used to filter out SNPs with any missing information, MAF <0.05, meanDP <5, or meanDP>80. Accession information for all raw RNA sequencing data used is available in Table 1.

### Population genomic analysis

Estimates for nucleotide diversity (π), between population divergence (*d*_XY_), density of fixed differences (*d*_f_), between population difference in allele frequency (*F*_ST_), and Tajima’s D were computed in non-overlapping sliding windows sized 10kb, 25kb, and 50kb, respectively. If a data set has substantial missing points, the estimation of nucleotide diversity is often biased (*80*). Thus, calculation of nucleotide diversity and *d*_XY_ from the variant call format (VCF) file can be over-estimated since missing sites are not distinguished from invariant (monomorphic) positions in variants-only VCFs. To account for potential inflation of population summary statistics, we considered an average breadth of coverage across samples for each window when we computed nucleotide diversity, *d*_XY_, and *d*_f_. Nucleotide diversity of protein coding sequence was computed using SNPGenie (*81*).

For phylogenetic reconstruction of ZAL2^m^ outlier windows, we used RaxML (ver 8.0.2) for 200kb windows. The targeted window was selected based on the Tajima’s D values and haplogroup assignments were determined by manually inspecting genotype plots. For the control region, we concatenated 10 randomly selected regions of 20kb windows (200kb). The evolutionary model was set to GTRGAMMA with acquisition bias correction using conditional likelihood method (-m ASC_GTRGAMMA –asc-corr=lewis). The medium ground finch (*Geospiza fortis*) was used as an outgroup species. We conducted bootstrap with 100 replicates.

The analysis of linkage disequilibrium (LD) decay was performed across the entire chromosomal rearrangement. We calculated pairwise *r*^2^ values between variant sites using *PopLDdecay* (*82*).

We performed H-scan (*63*) to identify soft and hard sweeps, inferred by extended tracts of homozygosity, using phased haplotypes of ZAL2 and ZAL2^m^. H-scan outputs were separated into 25kb non-overlapping windows. As a representative summary statistic, we chose the maximum H-statistic value for each window. To calculate the empirical *p*-value, we first binned genomic windows into 50 SNP increments based on the number of SNPs. When a window included more than 100 SNPs, all the windows were merged into one bin. On the basis of the ranking of summary statistics in each bin, we calculated an empirical *p*-value for each window. Candidate regions of positive selection were defined as those with an empirical *p*-value less than 0.05.

### Gene expression analyses

We examined both gene expression divergence and allele-specific expression using three RNAseq data sets from white-throated sparrows (four types of tissue from four separate samples of birds, see Table 1). The first data set consisted of gene expression in male white-throated sparrows collected during the breeding season. In those birds, the samples comprised two brain regions, the hypothalamus (Hyp) and the ventromedial arcopallium (AMV) (*25, 56*), a functional homolog of the medial amygdala (*83*). The second data set consisted of gene expression in the same two brain regions (Hyp and AMV) in adult females collected during the breeding season as well as male and female nestlings (see (*39*) for details). The third data set consisted of gene expression in heart and liver tissue in white-striped males collected during migration, then housed in captivity under two lighting conditions (long days and short days) and sampled at two time points during the day (ZT6 and ZT18) (*84*). In statistical analyses, the females and nestlings of each sex were treated as separate “batches” of RNAseq such that there were five total batches: brain samples from adult males, adult females, nestling males, nestling females, and liver/heart samples from adult males. Each tissue was nested within batch to account for repeated sampling from the same individuals.

RNAseq reads were aligned to a reference genome *N*-masked for putative fixed differences for both chromosomal polymorphisms present in this species (*i*.*e*., ZAL2/2, ZAL3^a^/3^a^,) using *HiSat2* (*85*). To examine allele-specific expression, we used *SNPsplit* (v0.3.2) to assign mapped reads to ZAL2 or ZAL2^m^ for the white-striped samples. Transcripts were quantified using *StringTie* (*86*) and differential expression and allelic-specific expression analyses were performed using *DESeq2* (*87*). To test for differential expression between the morphs, we used the following model in *DESeq2*: ‘design = ∼ morph’. To test for allelic bias (AB), we used the size factors generated for each sample in the previous step, and then used the following model: ‘design = ∼ individual + allele’. To perform hypothesis testing, we used linear mixed models with tissue nested in sequencing batch as random effects using the R package *lme4*. To test for the effect of interest, we then performed a likelihood ratio test to compare a full model to a reduced model with the factor of interest removed using the *anova* function. Where applicable, we used the *summary* function to perform post-hoc comparisons between multiple groups.

To test for significant allelic bias in expression within each sample type, we used a Wilcoxon rank sum test to test whether the ratio of ZAL2^m^ to ZAL2 expression differed significantly from 0.5, applying an FDR correction for multiple testing. To test for an association between allelic bias and the number of mutations within a gene, we computed the per-base number of mutations by dividing the number of mutations by the gene length and then sorted the genes into deciles. We also computed the number of fixed mutations within 1 kB upstream of the transcription start site (TSS). We then tested whether there was an association between allelic bias and the decile rank or number of mutations within 1 kB upstream of the TSS using the following linear model: Log2 AB ∼ Variable + (1|Batch:Tissue). To test for overrepresentation of morph-biased genes located inside the rearrangement, we performed a two-sample test for equality of proportions for each brain sample type only. For this test, we compared the proportion of differentially expressed genes, out of the 1007 genes inside the rearranged region on ZAL2, to the proportion out of the 13,369 genes located elsewhere in the genome. For this comparison, we used the *prop*.*test* function in R to perform a two-proportions Z-test, applying an FDR correction for multiple testing. To test whether the morph bias in expression of a gene significantly predicted allelic bias for that gene, we grouped genes into three categories: those that were more highly expressed in the white-striped morph (W>T), those more highly expressed in the tan-striped morph (T>W) and those that that did not differ significantly between morphs (T=W). Using the allelic bias for each gene, we then tested whether allelic bias differed among these categories using the following linear mixed model: Log2 AB ∼ MorphBias_Category + (1|Batch:Tissue), followed by Bonferroni-corrected pairwise post-hoc tests using the ‘ghlt’ function in *multcomp* package. Note that heart and liver tissue samples were obtained from white-striped birds; because we did not have data on differential expression by morph for these tissues, they were excluded from this analysis, as well as the tests for overrepresentation described above.

To test whether haplogroup affects gene expression, we merged the gene count matrices for the male (H1: *n* = 7, H2: *n* = 3) and female (H1: *n* = 3, H2: *n* = 3) white-striped birds that were also included in the whole genome sequencing dataset and tested for an effect of haplogroup using the following model: design ∼ Sex + Haplogroup. Only one gene in Hyp (Geranylgeranyl Diphosphate Synthase 1, *GGPS1*) and one gene in AMV (RUN And FYVE Domain Containing 4, *RUFY4*) were differentially expressed at the genome-wide level. Neither of these genes are located in the ZAL2^m^ outlier region and only *GGPS1* is located on ZAL2. Thus, we report only unadjusted *p*-values for genes inside the ZAL2^m^ outlier region that were differentially expressed (both unadjusted and adjusted *p*-values for all genes in the ZAL2m outlier region are reported in Supp. Table 4).

To examine the relationship between the H-statistic and allelic bias in expression, we computed the average H-statistic for each gene. Each gene was assigned the H-statistic value of the 20kb bin overlapping the gene (or the average of multiple 20kb bins, if a gene overlapped two bins). We then used a multiple linear regression model to examine the relationship between the Log_2_ASE and the log_2_H-statistic of a gene for both ZAL2 and ZAL2^m^ separately, using the following model: Log2 AB ∼ Log_2_H-statistic + (1|Batch:Tissue). Additionally, we performed functional enrichment analyses for the genes in regions with a significant H-statistic using *ToppFun* with human homolog HGNC gene names (*88*).

## Author Contributions

This sequencing effort and study design was conceived by DS, DLM and SVY. CNB was also involved in the early design of the study and was responsible for the acquisition of tissue samples from the Field Museum. DS, PC, TL were responsible for sample collection, generating sequencing libraries, and performing sequencing. HJ and DS were responsible for data curation and management. HJ, with assistance from DS, performed the genome assembly and population genomic analyses. NMB performed the RNA-sequencing analyses. Interpretation and writing were performed by HJ, NMB, DLM and SVY.

## Acknowledgements

We thank the members of the Maney lab who performed field work and collected the samples from GA and ME, as well as provided helpful feedback throughout the preparation of this manuscript. K.E. Grogan performed the RNA-sequencing for the samples from adult females and nestlings. David Willard (Collection Manager—Birds, Field Museum of Natural History, Chicago, IL) collected and provided access to the samples from IL. CNB thanks Dr. Elaina Tuttle for passing these samples on to him, and for introducing him to the white-throated sparrow research. This work was supported by NSF IOS-0723805 to DLM and by NIH 1R01MH082833, NIH R21MH102677, and NSF IOS-1656247 to DLM and SVY.

## References

1. D. Charlesworth, The status of supergenes in the 21st century: recombination suppression in Batesian mimicry and sex chromosomes and other complex adaptations. Evol Appl. 9, 74–90 (2016).

2. M. J. Thompson, C. D. Jiggins, Supergenes and their role in evolution. Heredity. 113, 1–8 (2014).

3. S. P. Otto, T. Lenormand, Resolving the paradox of sex and recombination. Nat Rev Genet. 3, 252– 261 (2002).

4. J. Wang, Y. Wurm, M. Nipitwattanaphon, O. Riba-Grognuz, Y.-C. Huang, D. Shoemaker, L. Keller, A Y-like social chromosome causes alternative colony organization in fire ants. Nature. 493, 664– 668 (2013).

5. Y.-C. Huang, V. D. Dang, N.-C. Chang, J. Wang, Multiple large inversions and breakpoint rewiring of gene expression in the evolution of the fire ant social supergene. P Roy Soc B-Biol Sci. 285, 20180221 (2018).

6. Z. Yan, S. H. Martin, D. Gotzek, S. V. Arsenault, P. Duchen, Q. Helleu, O. Riba-Grognuz, B. G. Hunt, N. Salamin, D. Shoemaker, K. G. Ross, L. Keller, Evolution of a supergene that regulates a trans-species social polymorphism. Nat Ecol Evol. 4, 240–249 (2020).

7. C. Martinez-Ruiz, R. Pracana, E. Stolle, C. I. Paris, R. A. Nichols, Y. Wurm, Genomic architecture and evolutionary antagonism drive allelic expression bias in the social supergene of red fire ants. eLife. 9, e55862 (2020).

8. L. L. Farrell, T. Burke, J. Slate, S. B. McRae, D. B. Lank, Genetic mapping of the female mimic morph locus in the ruff. BMC Genet. 14, 109 (2013).

9. C. Küpper, M. Stocks, J. E. Risse, N. dos Remedios, L. L. Farrell, S. B. McRae, T. C. Morgan, N. Karlionova, P. Pinchuk, Y. I. Verkuil, A. S. Kitaysky, J. C. Wingfield, T. Piersma, K. Zeng, J. Slate, M. Blaxter, D. B. Lank, T. Burke, A supergene determines highly divergent male reproductive morphs in the ruff. Nat Genet. 48, 79–83 (2016).

10. S. Lamichhaney, G. Fan, F. Widemo, U. Gunnarsson, D. S. Thalmann, M. P. Hoeppner, S. Kerje, U. Gustafson, C. Shi, H. Zhang, W. Chen, X. Liang, L. Huang, J. Wang, E. Liang, Q. Wu, S. M.-Y. Lee, X. Xu, J. Höglund, X. Liu, L. Andersson, Structural genomic changes underlie alternative reproductive strategies in the ruff (Philomachus pugnax). Nat Genet. 48, 84–88 (2016).

11. T. Schwander, R. Libbrecht, L. Keller, Supergenes and complex phenotypes. Curr Biol. 24, R288– R294 (2014).

12. D. E. Pearse, N. J. Barson, T. Nome, G. Gao, M. A. Campbell, A. Abadía-Cardoso, E. C. Anderson, D. E. Rundio, T. H. Williams, K. A. Naish, T. Moen, S. Liu, M. Kent, M. Moser, D. R. Minkley, E. B. Rondeau, M. S. O. Brieuc, S. R. Sandve, M. R. Miller, L. Cedillo, K. Baruch, A. G. Hernandez, G. Ben-Zvi, D. Shem-Tov, O. Barad, K. Kuzishchin, J. C. Garza, S. T. Lindley, B. F. Koop, G. H. Thorgaard, Y. Palti, S. Lien, Sex-dependent dominance maintains migration supergene in rainbow trout. Nat Ecol Evol. 3, 1731–1742 (2019).

13. E. R. Hager, O. S. Harringmeyer, T. B. Wooldridge, S. Theingi, J. T. Gable, S. McFadden, B. Neugeboren, K. M. Turner, H. E. Hoekstra, bioRxiv (2021) doi:10.1101/2021.01.21.427490..

14. M. Joron, L. Frezal, R. T. Jones, N. L. Chamberlain, S. F. Lee, C. R. Haag, A. Whibley, M. Becuwe, S. W. Baxter, L. Ferguson, P. A. Wilkinson, C. Salazar, C. Davidson, R. Clark, M. A. Quail, H. Beasley, R. Glithero, C. Lloyd, S. Sims, M. C. Jones, J. Rogers, C. D. Jiggins, R. H. ffrench-Constant, Chromosomal rearrangements maintain a polymorphic supergene controlling butterfly mimicry. Nature. 477, 203–206 (2011).

15. K. Kunte, W. Zhang, A. Tenger-Trolander, D. H. Palmer, A. Martin, R. D. Reed, S. P. Mullen, M. R. Kronforst, doublesex is a mimicry supergene. Nature. 507, 229–232 (2014).

16. T. Kess, P. Bentzen, S. J. Lehnert, E. V. A. Sylvester, S. Lien, M. P. Kent, M. Sinclair-Waters, C. J. Morris, P. Regular, R. Fairweather, I. R. Bradbury, A migration-associated supergene reveals loss of biocomplexity in Atlantic cod. Sci Adv. 5, eaav2461 (2019).

17. M. Lundberg, M. Liedvogel, K. Larson, H. Sigeman, M. Grahn, A. Wright, S. Åkesson, S. Bensch, Genetic differences between willow warbler migratory phenotypes are few and cluster in large haplotype blocks. Evol Lett. 1, 155–168 (2017).

18. R. B. Roberts, J. R. Ser, T. D. Kocher, Sexual conflict resolved by invasion of a novel sex determiner in Lake Malawi cichlid fishes. Science. 326, 998–1001 (2009).

19. I. Sanchez-Donoso, S. Ravagni, J.D. Rodríguez-Teijeiro, M. J. Christmas, Y. Huang, A. Maldonado-Linares, M. Puigcerver, I. Jiménez-Blasco, P. Andrade, D. Gonçalves, G. Friis, I. Roig, M. T. Webster, J. A. Leonard, C. Vilà, Massive genome inversion drives coexistence of divergent morphs in common quails. Curr Biol. 32, 462-469.e6 (2022).

20. E. R. Funk, N. A. Mason, S. Pálsson, T. Albrecht, J. A. Johnson, S. A. Taylor, A supergene underlies linked variation in color and morphology in a Holarctic songbird. Nat Commun. 12, 6833 (2021).

21. E. M. Tuttle, Alternative reproductive strategies in the white-throated sparrow: behavioral and genetic evidence. Behav. Ecol. 14, 425–432 (2003).

22. D. L. Maney, B. M. Horton, W. M. Zinzow-Kramer, Estrogen receptor alpha as a mediator of life-history trade-offs. Integr. Comp. Biol. 55, 323–331 (2015).

23. B. M. Horton, Y. Hu, C. L. Martin, B. P. Bunke, B. S. Matthews, I. T. Moore, J. W. Thomas, D. L. Maney, Behavioral characterization of a white-throated sparrow homozygous for the ZAL2m chromosomal rearrangement. Behav Genet. 43, 60–70 (2013).

24. E. M. Tuttle, A. O. Bergland, M. L. Korody, M. S. Brewer, D. J. Newhouse, P. Minx, M. Stager, A. Betuel, Z. A. Cheviron, W. C. Warren, R. A. Gonser, C. N. Balakrishnan, Divergence and functional degradation of a sex chromosome-like supergene. Curr. Biol. 26, 344–350 (2016).

25. D. Sun, I. Huh, W. M. Zinzow-Kramer, D. L. Maney, S. V. Yi, Rapid regulatory evolution of a nonrecombining autosome linked to divergent behavioral phenotypes. PNAS. 115, 2794–2799 (2018).

26. J. R. Merritt, K. E. Grogan, W. M. Zinzow-Kramer, D. Sun, E. A. Ortlund, S. V. Yi, D. L. Maney, A supergene-linked estrogen receptor drives alternative phenotypes in a polymorphic songbird. PNAS (2020), doi:10.1073/pnas.2011347117..

27. J. K. Lowther, Polymorphism in the white-throated sparrow, Zonotrichia albicollis (Gmelin). Can. J. Zool. 39, 281–292 (1961).

28. D. L. Maney, Endocrine and genomic architecture of life history trade-offs in an avian model of social behavior. Gen. Comp. Endocrinol. 157, 275–282 (2008).

29. B. M. Horton, M. E. Hauber, D. L. Maney, Morph matters: Aggression bias in a polymorphic sparrow. PLOS ONE. 7, e48705 (2012).

30. H. B. Thorneycroft, Chromosomal polymorphism in the white-throated sparrow, Zonotrichia albicollis (Gmelin). Science. 154, 1571–1572 (1966).

31. H. B. Thorneycroft, A cytogenetic study of the white-throated sparrow, Zonotrichia albicollis (Gmelin). Evolution. 29, 611–621 (1975).

32. J. W. Thomas, M. Cáceres, J. J. Lowman, C. B. Morehouse, M. E. Short, E. L. Baldwin, D. L. Maney, C. L. Martin, The chromosomal polymorphism linked to variation in social behavior in the white-throated sparrow (Zonotrichia albicollis) Is a complex rearrangement and suppressor of recombination. Genetics. 179, 1455–1468 (2008).

33. L. Y. Huynh, D. L. Maney, J. W. Thomas, Contrasting population genetic patterns within the white-throated sparrow genome (Zonotrichia albicollis). BMC Genet. 11, 96 (2010).

34. L. Campagna, Supergenes: The genomic architecture of a bird with four sexes. Curr Biol. 26, R105– R107 (2016).

35. J. B. Falls, J. G. Kopachena, in Birds of the World, J. F. Poole, Ed. (Cornell Lab of Ornithology, Ithaca, NY, 2020; https://doi.org/10.2173/bow.whtspa.01).

36. N. H. Barton, B. Charlesworth, Why sex and recombination? Science (1998), doi:10.1126/science.281.5385.1986..

37. B. Charlesworth, The effects of deleterious mutations on evolution at linked sites. Genetics. 190, 5–22 (2012).

38. D. L. Maney, J. R. Merritt, M. R. Prichard, B. M. Horton, S. V. Yi, Inside the supergene of the bird with four sexes. Horm Behav. 126, 104850 (2020).

39. D. Sun, T. S. Layman, H. Jeong, P. Chatterjee, K. Grogan, J. R. Merritt, D. L. Maney, S. V. Yi, Genome-wide variation in DNA methylation linked to developmental stage and chromosomal suppression of recombination in white-throated sparrows. Mol Ecol. 30, 3453–3467 (2021).

40. J. K. Davis, L. B. Mittel, J. J. Lowman, P. J. Thomas, D. L. Maney, C. L. Martin, J. W. Thomas, Haplotype-based genomic sequencing of a chromosomal polymorphism in the white-throated sparrow (Zonotrichia albicollis). J Hered. 102, 380–390 (2011).

41. R. A. Fisher, The evolution of dominance. Biol Rev. 6, 345–368 (1931).

42. J. J. Bull, Evolution of Sex Determining Mechanisms (Benjamin-Cummings Pub Co, London, 1983).

43. D. Charlesworth, B. Charlesworth, Sex differences in fitness and selection for centric fusions between sex-chromosomes and autosomes. Genet Res. 35, 205–214 (1980).

44. W. R. Rice, The accumulation of sexually antagonistic genes as a selective agent promoting the evolution of reduced recombination between primitive sex chromosomes. Evolution. 41, 911–914 (1987).

45. W. R. Rice, Genetic hitchhiking and the evolution of reduced genetic activity of the Y sex chromosome. Genetics. 116, 161–167 (1987).

46. B. M. Horton, C. M. Michael, M. R. Prichard, D. L. Maney, Vasoactive intestinal peptide as a mediator of the effects of a supergene on social behaviour. P Roy Soc B-Biol Sci. 287, 20200196 (2020).

47. J. L. Goodson, A. M. Kelly, M. A. Kingsbury, R. R. Thompson, An aggression-specific cell type in the anterior hypothalamus of finches. PNAS. 109, 13847–13852 (2012).

48. D. Bachtrog, Evidence that positive selection drives Y-chromosome degeneration in Drosophila miranda. Nat Genet. 36, 518–522 (2004).

49. D. Bachtrog, A dynamic view of sex chromosome evolution. Curr Opin Genet Dev. 16, 578–585 (2006).

50. Q. Zhou, D. Bachtrog, Sex-specific adaptation drives early sex chromosome evolution in Drosophila. Science. 337, 341–345 (2012).

51. S. Glémin, Balancing selection in self-fertilizing populations. Evolution. 75, 1011–1029 (2021).

52. S. Glémin, C. M. François, N. Galtier, in Evolutionary Genomics: Statistical and Computational Methods, M. Anisimova, Ed. (Springer, New York, NY, 2019; https://doi.org/10.1007/978-1-4939-9074-0_11), Methods in Molecular Biology, pp. 331–369.

53. L. F. Delph, J. K. Kelly, On the importance of balancing selection in plants. New Phytol. 201, 45– 56 (2014).

54. B. S. Gaut, C. M. Díez, P. L. Morrell, Genomics and the contrasting dynamics of annual and perennial domestication. Trends Genet. 31, 709–719 (2015).

55. K. M. Siewert, B. F. Voight, BetaScan2: Standardized statistics to detect balancing selection utilizing substitution data. Genome Biol Evol. 12, 3873–3877 (2020).

56. W. M. Zinzow-Kramer, B. M. Horton, C. D. McKee, J. M. Michaud, G. K. Tharp, J. W. Thomas, E. M. Tuttle, S. Yi, D. L. Maney, Genes located in a chromosomal inversion are correlated with territorial song in white-throated sparrows. Genes Brain Behav. 14, 641–654 (2015).

57. B. M. Horton, W. H. Hudson, E. A. Ortlund, S. Shirk, J. W. Thomas, E. R. Young, W. M. Zinzow-Kramer, D. L. Maney, Estrogen receptor α polymorphism in a species with alternative behavioral phenotypes. PNAS. 111, 1443–1448 (2014).

58. D. Georgiev, D. Arion, J. F. Enwright, M. Kikuchi, Y. Minabe, J. P. Corradi, D. A. Lewis, T. Hashimoto, Lower gene expression for KCNS3 potassium channel subunit in parvalbumin-containing neurons in the prefrontal cortex in schizophrenia. AJP. 171, 62–71 (2014).

59. T. Miyamae, T. Hashimoto, M. Abraham, R. Kawabata, S. Koshikizawa, Y. Bian, Y. Nishihata, M. Kikuchi, G. B. Ermentrout, D. A. Lewis, G. Gonzalez-Burgos, Kcns3 deficiency disrupts Parvalbumin neuron physiology in mouse prefrontal cortex: Implications for the pathophysiology of schizophrenia. Neurobiol Dis. 155, 105382 (2021).

60. B. T. Lahn, D. C. Page, Four evolutionary strata on the human X chromosome. Science. 286, 964– 967 (1999).

61. L. Xu, G. Auer, V. Peona, A. Suh, Y. Deng, S. Feng, G. Zhang, M. P. K. Blom, L. Christidis, S. Prost, M. Irestedt, Q. Zhou, Dynamic evolutionary history and gene content of sex chromosomes across diverse songbirds. Nat Ecol Evol, 1 (2019).

62. R. Bergero, A. Forrest, E. Kamau, D. Charlesworth, Evolutionary strata on the X chromosomes of the dioecious plant Silene latifolia: Evidence from new sex-linked genes. Genetics. 175, 1945–1954 (2007).

63. Messer Lab — Resources. Messer Lab (2014), (available at https://messerlab.org/resources/).

64. B. Charlesworth, P. H. Harvey, B. Charlesworth, D. Charlesworth, The degeneration of Y chromosomes. Phil Trans R Soc B. 355, 1563–1572 (2000).

65. S. Yi, B. Charlesworth, Contrasting patterns of molecular evolution of the genes on the new and old sex chromosomes of Drosophila miranda. Mol Biol Evol. 17, 703–717 (2000).

66. D. Bachtrog, B. Charlesworth, Reduced adaptation of a non-recombining neo-Y chromosome. Nature. 416, 323–326 (2002).

67. N. D. Singh, L. B. Koerich, A. B. Carvalho, A. G. Clark, Positive and purifying selection on the Drosophila Y chromosome. Mol Biol Evol. 31, 2612–2623 (2014).

68. W. R. Rice, Sex chromosomes and the evolution of sexual dimorphism. Evolution. 38, 735–742 (1984).

69. B. Vicoso, B. Charlesworth, Evolution on the X chromosome: unusual patterns and processes. Nat Rev Genet. 7, 645–653 (2006).

70. J. E. Mank, Small but mighty: the evolutionary dynamics of W and Y sex chromosomes. Chromosome Res. 20, 21–33 (2012).

71. A. L. Evans, P. A. Mena, B. F. McAllister, Positive selection near an inversion breakpoint on the neo-X chromosome of Drosophila americana. Genetics. 177, 1303–1319 (2007).

72. K. Nam, K. Munch, A. Hobolth, J. Y. Dutheil, K. R. Veeramah, A. E. Woerner, M. F. Hammer, G. A. G. D. Project, T. Mailund, M. H. Schierup, Extreme selective sweeps independently targeted the X chromosomes of the great apes. PNAS. 112, 6413–6418 (2015).

73. B. Gegenhuber, M. V. Wu, R. Bronstein, J. Tollkuhn, Regulation of neural gene expression by estrogen receptor alpha, bioRxiv (2020), doi:10.1101/2020.10.21.349290..

74. M. Wellenreuther, L. Bernatchez, Eco-evolutionary genomics of chromosomal inversions. Trends in Ecology & Evolution. 33, 427–440 (2018).

75. C. N. Balakrishnan, M. Mukai, R. A. Gonser, J. C. Wingfield, S. E. London, E. M. Tuttle, D. F. Clayton, Brain transcriptome sequencing and assembly of three songbird model systems for the study of social behavior. PeerJ. 2, e396 (2014).

76. A. M. Bolger, M. Lohse, B. Usadel, Trimmomatic: a flexible trimmer for Illumina sequence data. Bioinformatics. 30, 2114–2120 (2014).

77. L. Song, L. Florea, B. Langmead, Lighter: fast and memory-efficient sequencing error correction without counting. Genome Biol. 15, 509 (2014).

78. S. D. Jackman, B. P. Vandervalk, H. Mohamadi, J. Chu, S. Yeo, S. A. Hammond, G. Jahesh, H. Khan, L. Coombe, R. L. Warren, I. Birol, ABySS 2.0: resource-efficient assembly of large genomes using a Bloom filter. Genome Res. 27, 768–777 (2017).

79. Y. Xie, G. Wu, J. Tang, R. Luo, J. Patterson, S. Liu, W. Huang, G. He, S. Gu, S. Li, X. Zhou, T.-W. Lam, Y. Li, X. Xu, G. K.-S. Wong, J. Wang, SOAPdenovo-Trans: de novo transcriptome assembly with short RNA-Seq reads. Bioinformatics. 30, 1660–1666 (2014).

80. K. L. Korunes, K. Samuk, pixy: Unbiased estimation of nucleotide diversity and divergence in the presence of missing data. Mol Ecol Resour. 21, 1359–1368 (2021).

81. C. W. Nelson, L. H. Moncla, A. L. Hughes, SNPGenie: estimating evolutionary parameters to detect natural selection using pooled next-generation sequencing data. Bioinformatics. 31, 3709–3711 (2015).

82. C. Zhang, S.-S. Dong, J.-Y. Xu, W.-M. He, T.-L. Yang, PopLDdecay: a fast and effective tool for linkage disequilibrium decay analysis based on variant call format files. Bioinformatics. 35, 1786– 1788 (2019).

83. C. V. Mello, T. Kaser, A. A. Buckner, M. Wirthlin, P. V. Lovell, Molecular architecture of the zebra finch arcopallium. J Comp Neurol. 527, 2512–2556 (2019).

84. W. J. Horton, M. Jensen, A. Sebastian, C. A. Praul, I. Albert, P. A. Bartell, Transcriptome analyses of heart and liver reveal novel pathways for regulating songbird migration. Sci Rep. 9, 6058 (2019).

85. D. Kim, B. Langmead, S. L. Salzberg, HISAT: a fast spliced aligner with low memory requirements. Nat. Methods. 12, 357–360 (2015).

86. M. Pertea, D. Kim, G. M. Pertea, J. T. Leek, S. L. Salzberg, Transcript-level expression analysis of RNA-seq experiments with HISAT, StringTie and Ballgown. Nat Protoc. 11, 1650–1667 (2016).

87. M. I. Love, W. Huber, S. Anders, Moderated estimation of fold change and dispersion for RNA-seq data with DESeq2. Genome Biol. 15, 550 (2014).

88. J. Chen, E. E. Bardes, B. J. Aronow, A. G. Jegga, ToppGene Suite for gene list enrichment analysis and candidate gene prioritization. Nucleic Acids Res. 37, W305–W311 (2009).

