## Supplementary Materials for "Dynamic molecular evolution of a supergene with suppressed recombination in white-throated sparrows"

**Supplementary Figures:**

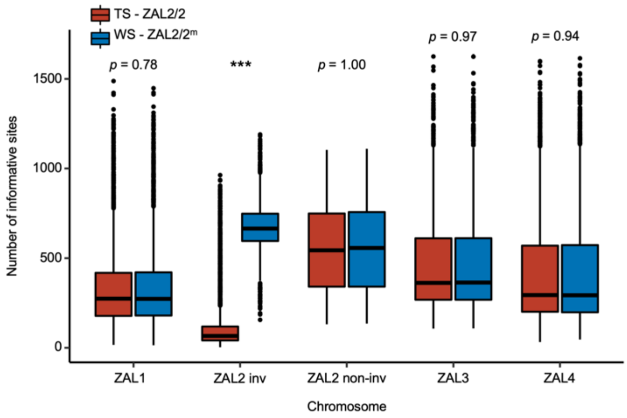


**Supplementary Fig. 1:** *The number of informative sites inside the ZAL2m rearrangement differed between morphs.* The number of informative sites in tan-striped versus white-striped birds is shown for the four largest chromosomes (macrochromosomes), computed using the same number of samples from tan-striped and white-striped birds ($N$ = 13 each). ZAL2 inv = inverted region in ZAL2 or ZAL2$^{m}$; ZAL2 non-inv = non-inverted region in ZAL2 or ZAL2$^{m}$.

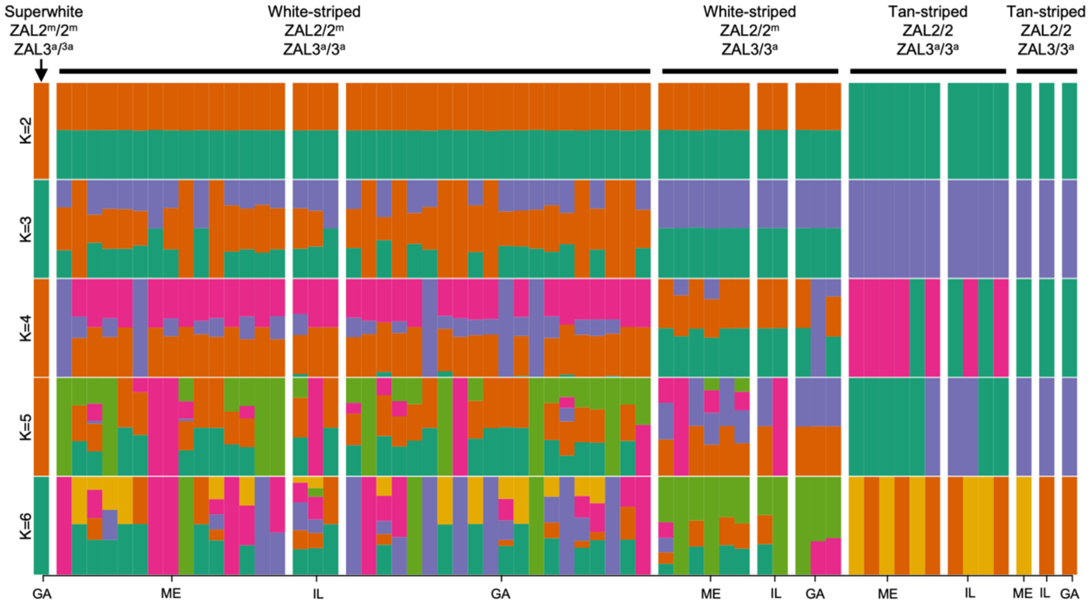


**Supplementary Fig. 2:** *Admixture tests showed no population substructure by geographic sampling location*. Inferred ancestral population fractions are shown for each bird as estimated by ADMIXTURE (K = 2 to K = 6 possible populations) for birds of each genotype and from different sampling locations. ADMIXTURE was run using all SNPs in the genome, excluding SNPs that met any of the following criteria: MAF < 0.01, missing > 20%, located inside the additional chromosomal polymorphism on ZAL3, or located in sex chromosomes. Note that 'geographic location' here refers to the site of collection or capture of the bird. Breeding locations for the GA and IL birds were unknown

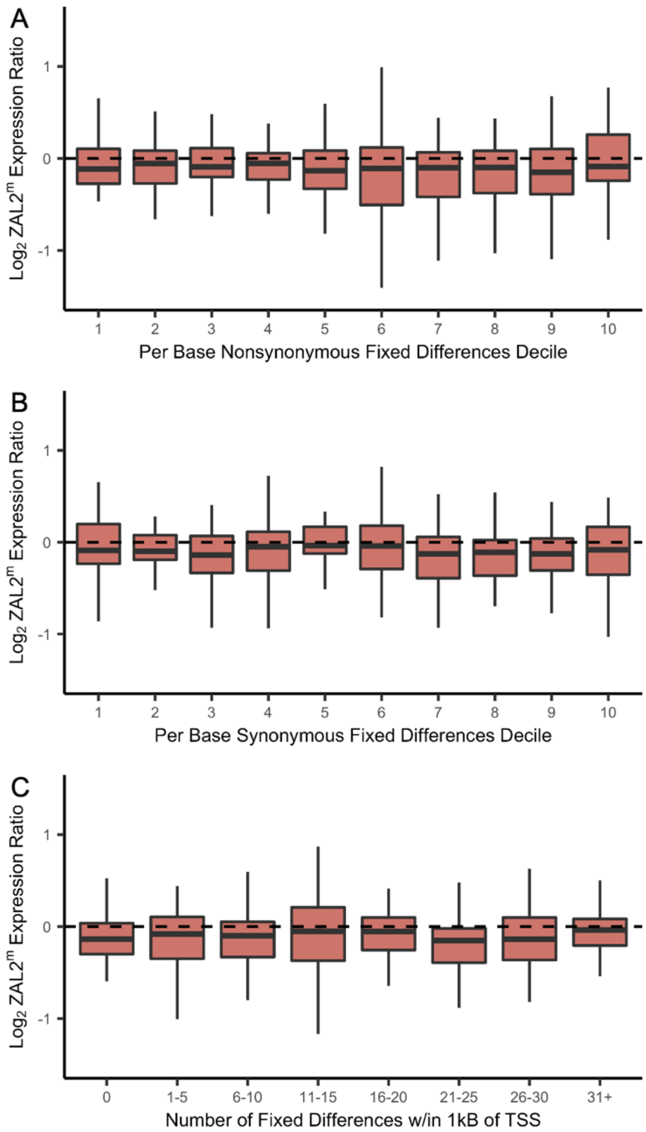


**Supplementary Figure 3.** *Allelic bias in expression was associated with the number of non-synonymous fixed differences.* Allelic bias in expression for each gene, averaged across sequencing batch and tissue (see Table 1, Materials & Methods), is plotted by the decile rank of the number of per-base fixed differences for that gene that are A) non-synonymous, B) synonymous, or C) the number of fixed differences within 1kb upstream of the transcription start site. Only the decile rank of the per-base number of non-synonymous fixed differences was associated with allelic bias in gene expression ($X^2(1) = 12.54$, $p = 0.000398$).

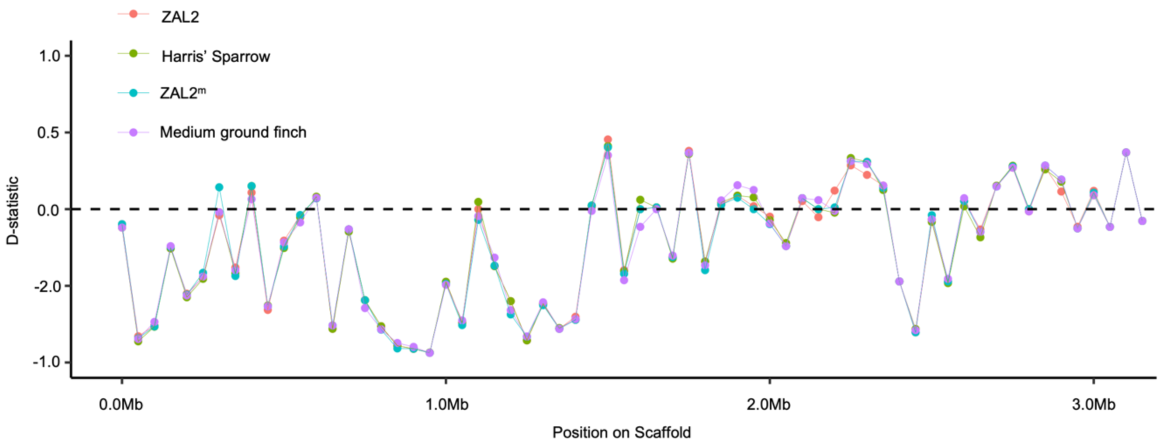


**Supplementary Fig. 4:** The *D-statistic did not vary by haplotype.* 50kb sliding window estimates of the D-statistic resulting from ABBA-BABA tests of ZAL2, Harris' sparrow (*Zonotrichia querula*), ZAL2$^{m}$, and Medium ground finch (*Geospiza fortis*) are plotted for four different haplotype groups in the ZAL2$^{m}$ outlier region.

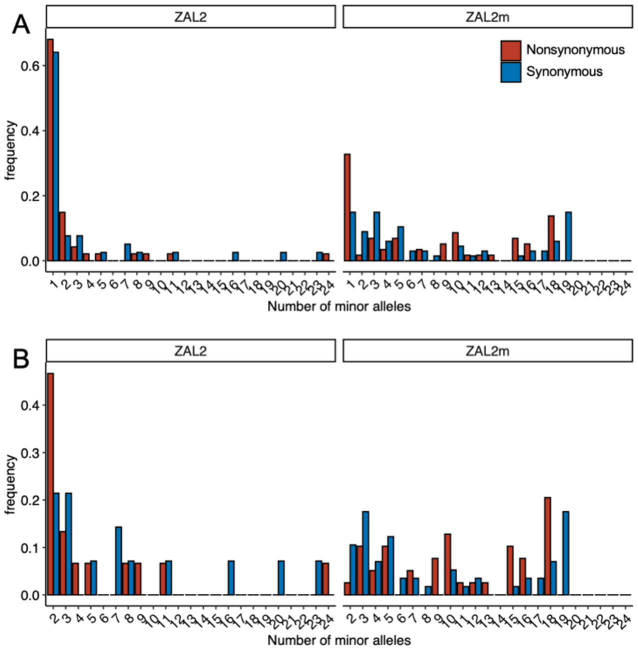


**Supplementary Fig. 5:** *The ZAL2$^{m}$ outlier region exhibited an excess of intermediate frequency minor alleles*. Site frequency spectra of polymorphic sites inside the ZAL2$^{m}$ outlier region are shown for both the ZAL2 and ZAL2$^{m}$ chromosomes. A) shows all ZAL2/ZAL2$^{m}$-linked SNPs and B) excludes singleton SNPs.

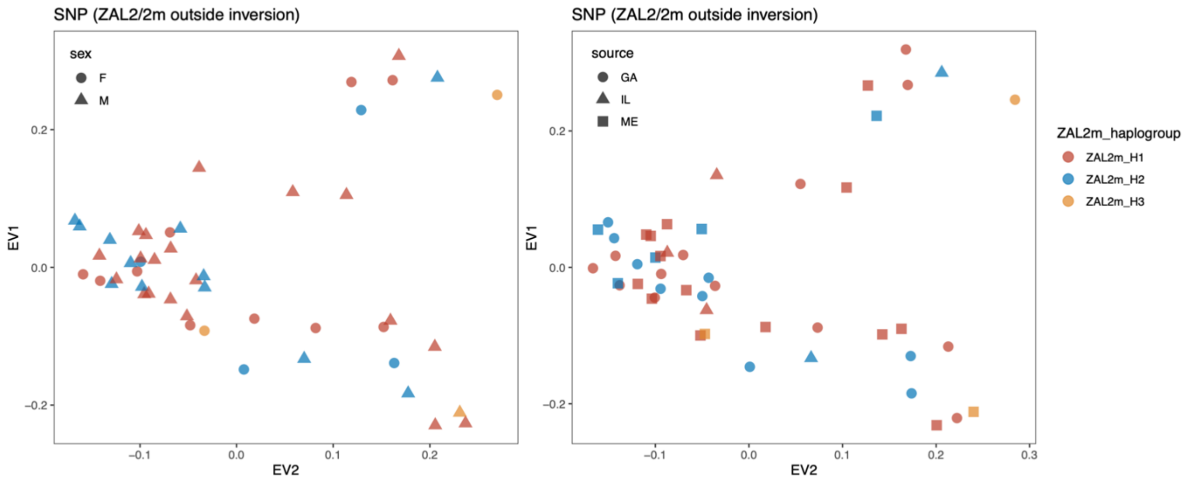


**Supplementary Fig. 6:** *Neither sex nor geographic location of sample collection produced distinct patterns between haplogroups*. Scatterplots of eigenvector 1 (PC1) and eigenvector 2 (PC2) from principal component analysis of all single-nucleotide variants (SNP) outside the ZAL2/2m inversion are shown. Colors show the haplogroup of the sample. In the left panel, shape indicates the sex of the sample and in the right panel, shape indicates the geographic sampling location. Note that the GA and IL birds were captured during migration, so their breeding location was unknown.

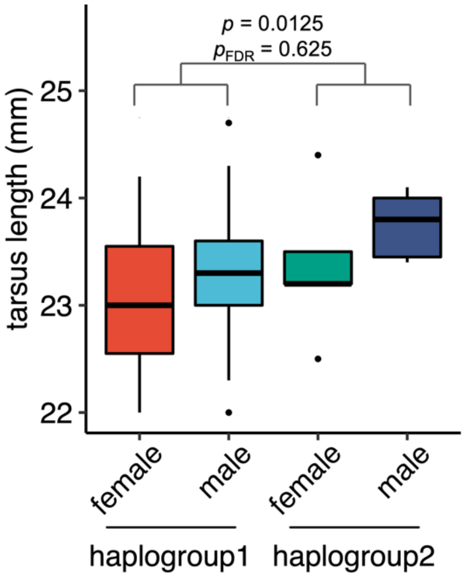


**Supplementary Fig. 7:** *Tarsus length did not differ by haplogroup in white-striped birds*. Boxplot of the tarsus length (mm) in birds of haplogroup 1 (# of female = 7, # of male = 15) and haplogroup 2 (# of female = 5, # of male = 6). We used a linear model to test the effect of haplogroup, while controlling for sex, using the following model: tarsus_length ~ sex + haplogroup. Although the effect was significant at $p < 0.05$, it did not survive FDR correction for multiple testing (50 tests were conducted). The marginal $r^2$ = 0.110 for the effect of haplogroup in the model, which is a medium effect size.

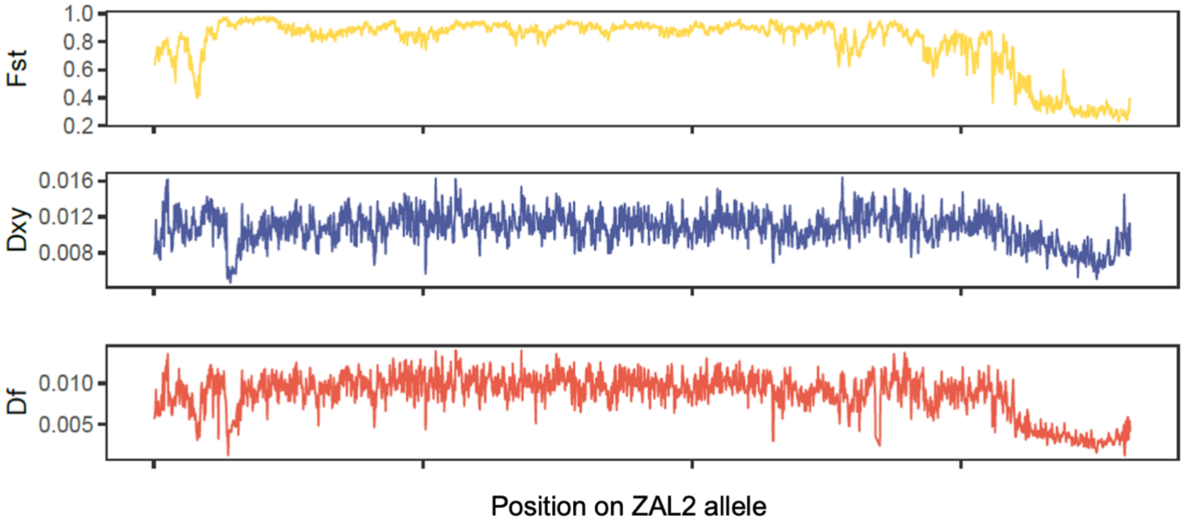


**Supplementary Fig. 8:** *Genetic differentiation between ZAL2 and ZAL2m is reduced at the ends of the chromosomal arms*. Plots show the population differentiation in allele frequency ($F_{ST}$) between tan and white birds, the number of nucleotide substitutions per site ($d_{XY}$) between ZAL2 and ZAL2$^m$, and density of fixed differences ($d_f$) between ZAL2 and ZAL2$^m$ inside the rearranged region.

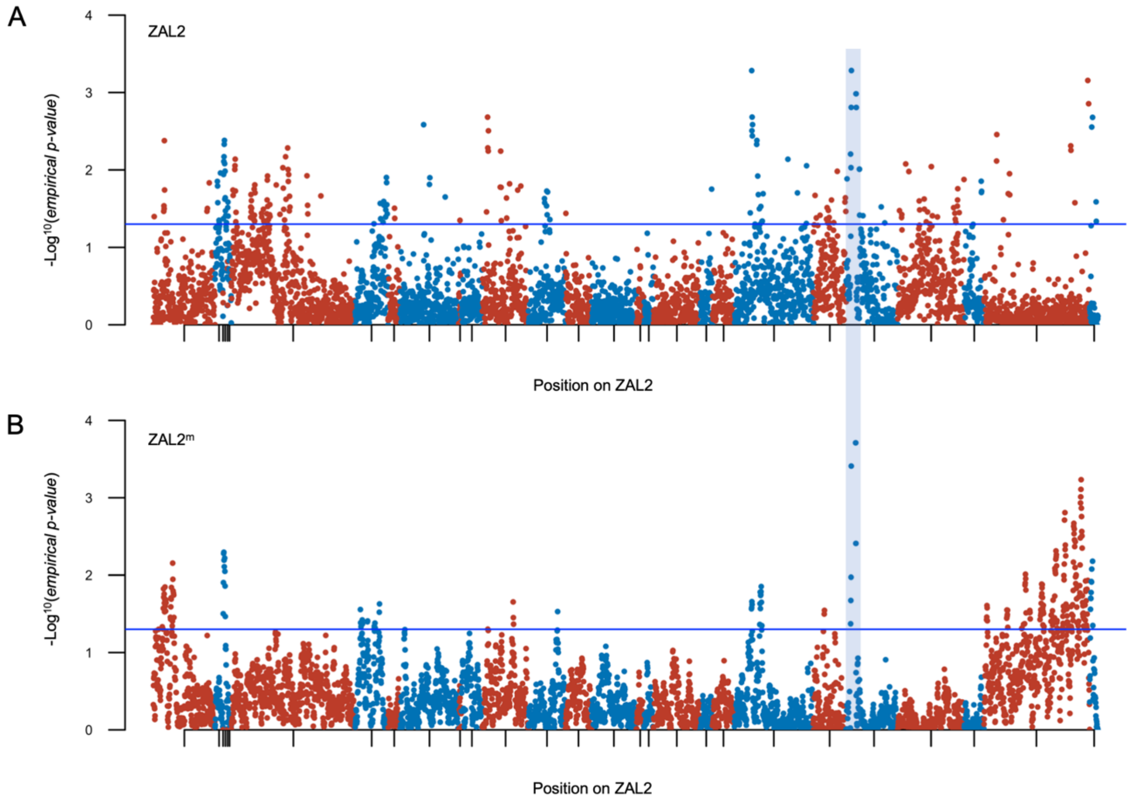


**Supplementary Figure 9.** *Both ZAL2 and ZAL2$^{m}$ have experienced selective sweeps*. The imputed *p*-values of the H-statistic (a measure of homozygosity, computed in 20kb windows) are plotted along the position on ZAL2 for A) ZAL2 and B) ZAL2$^{m}$. Colors refer to alternating scaffolds. A candidate region showing positive selection on both ZAL2 and ZAL2$^{m}$ (NW_005081582.1, 480-520kb and 920-960kb) is highlighted in blue.

**Supplementary Table 2:** *Population genetics sequencing information*

| | # birds | Total # reads | Total depth of coverage |
|---|---|---|---|
| WS | 49 | 14.9 B | 2,037x |
| TS | 13 | 2.7 B | 369x |

**Supplementary Table 3:** *Summary statistics of the genome assembly*

| | TS Assembly[1] | SWS assembly |
|---|---|---|
| Longest scaffold | 45,240,865 | 27,536,075 |
| Number of scaffolds > 1K nt | 5,673 | 14,669 |
| Number of scaffolds > 10K nt | 1,611 | 2,092 |
| Number of scaffolds > 100K nt | 535 | 672 |
| Number of scaffolds > 1M nt | 210 | 244 |
| Number of scaffolds > 10M nt | 18 | 13 |
| N50 scaffold length | 4,866,725 | 3,189,904 |
| L50 scaffold count | 52 | 81 |
| N50 contig length | 116,237 | 216,976 |
| L50 contig count | 2,295 | 1,331 |

[1] Tuttle et al., 2016
